# Ancestral Roles of the Fam20C Family of Secreted Protein Kinases Revealed by Functional Analysis in *C. elegans*

**DOI:** 10.1101/363440

**Authors:** Adina Gerson-Gurwitz, Carolyn A. Worby, Kian-Yong Lee, Renat Khaliullin, Jeff Bouffard, Dhanya Cheerambathur, Erin J. Cram, Karen Oegema, Jack E. Dixon, Arshad Desai

## Abstract

Fam20C is a secreted protein kinase mutated in Raine syndrome, a human skeletal disorder. In vertebrates, bone and enamel proteins are major Fam20C substrates. However, Fam20 kinases are conserved in invertebrates lacking bone and enamel, suggesting other ancestral functions. We show that FAMK-1, the *C. elegans* Fam20C ortholog, contributes to fertility, embryogenesis, and development. These functions are not fulfilled when FAMK-1 is retained in the early secretory pathway. During embryogenesis, FAMK-1 maintains inter-cellular partitions and prevents multinucleation; notably, temperature elevation or lowering cortical stiffness reduces requirement for FAMK-1 in this context. FAMK-1 is expressed in multiple adult tissues that undergo repeated mechanical strain, and selective expression in the spermatheca restores fertility. Informatic, biochemical and functional analysis implicate lectins as FAMK-1 substrates. These findings suggest that FAMK-1 phosphorylation of substrates, including lectins, in the late secretory pathway is important in embryonic and tissue contexts where cells are subjected to mechanical strain.

## INTRODUCTION

The recent discovery of kinases that reside in the secretory pathway and/or are secreted into the extracellular environment has sparked interest in the substrates of these kinases as well as the functional significance of extracellular phosphorylation. One family of secreted kinases, distantly related to the bacterial kinase, HipA, includes Fam20A, Fam20B, Fam20C, Fam198A, Fam198B, and *Drosophila* Four-jointed (Fj) (Tagliabracci et al. 2013; Sreelatha et al. 2015). Originally, these proteins were designated Fams because of shared sequence similarities and lack of information about their functions. These kinases reside in the oxidizing environment of the secretory pathway or outside the cell and they differ sufficiently from canonical kinases that they were not included in the human “kinome” (Manning et al. 2002). To date, these kinases either phosphorylate protein or sugar substrates. Of these, Fam20C was shown to be the authentic Golgi casein kinase, phosphorylating secreted proteins such as casein and the small integrin-binding ligand, N-linked glycoproteins (SIBLINGS) on SerXGlu/pSer (SXE/pS) motifs (Bahl et al. 2008; Zhou et al. 2009; Salvi et al. 2010; Tagliabracci et al. 2012). These motifs represent the majority of sites identified in secreted phospho-proteomes, and Fam20C was shown to generate the majority of the extracellular phosphoproteome in liver, breast epithelial and osteoblast-like cell lines, suggesting roles for Fam20C in a number of different biological processes (Cui et al. 2015; Tagliabracci et al. 2015).

Fam20C in mammals is most closely related to Fam20A and Fam20B. However, despite considerable sequence conservation, these proteins have very different functions. Fam20A is a pseudokinase that functions to activate Fam20C in specific tissues such as the lactating mammary gland and ameloblasts (Cui et al. 2015). Fam20B is not a protein kinase but instead is a xylose kinase involved in the maturation of proteoglycan chains (Koike et al. 2009; Wen et al. 2014). Work carried out in zebrafish demonstrates that Fam20B mutants fail to produce adequate levels of chondroitin sulfate proteoglycans (Eames et al. 2011). While vertebrates have all three members of the Fam20 family represented in their genomes, invertebrate organisms do not have a gene encoding Fam20A. *Drosophila melanogaster* contains both Fam20C- and Fam20B-like kinases but other invertebrates such as *C. elegans* (nematode) *C. intestinalis* (sea squirt) and *A. queenslandica* (marine sponge) contain only one family member, presumably representing either a Fam20C- or Fam20B-like activity (**Fig. 1 A**)(Tagliabracci et al. 2012). It was initially unclear whether the *C. elegans* Fam20-related protein has C-like and/or B-like activity, as it shared comparable sequence similarity with mammalian Fam20C (63%) and Fam20B (57%) in the atypical kinase domain (Xiao et al. 2013). Kinase assays utilizing a Fam20C-specific peptide or Fam20B-specific tetrasaccharide unequivocally demonstrated that *C. elegans* Fam20 functions as a protein kinase (Xiao et al. 2013).

**Figure 1.**
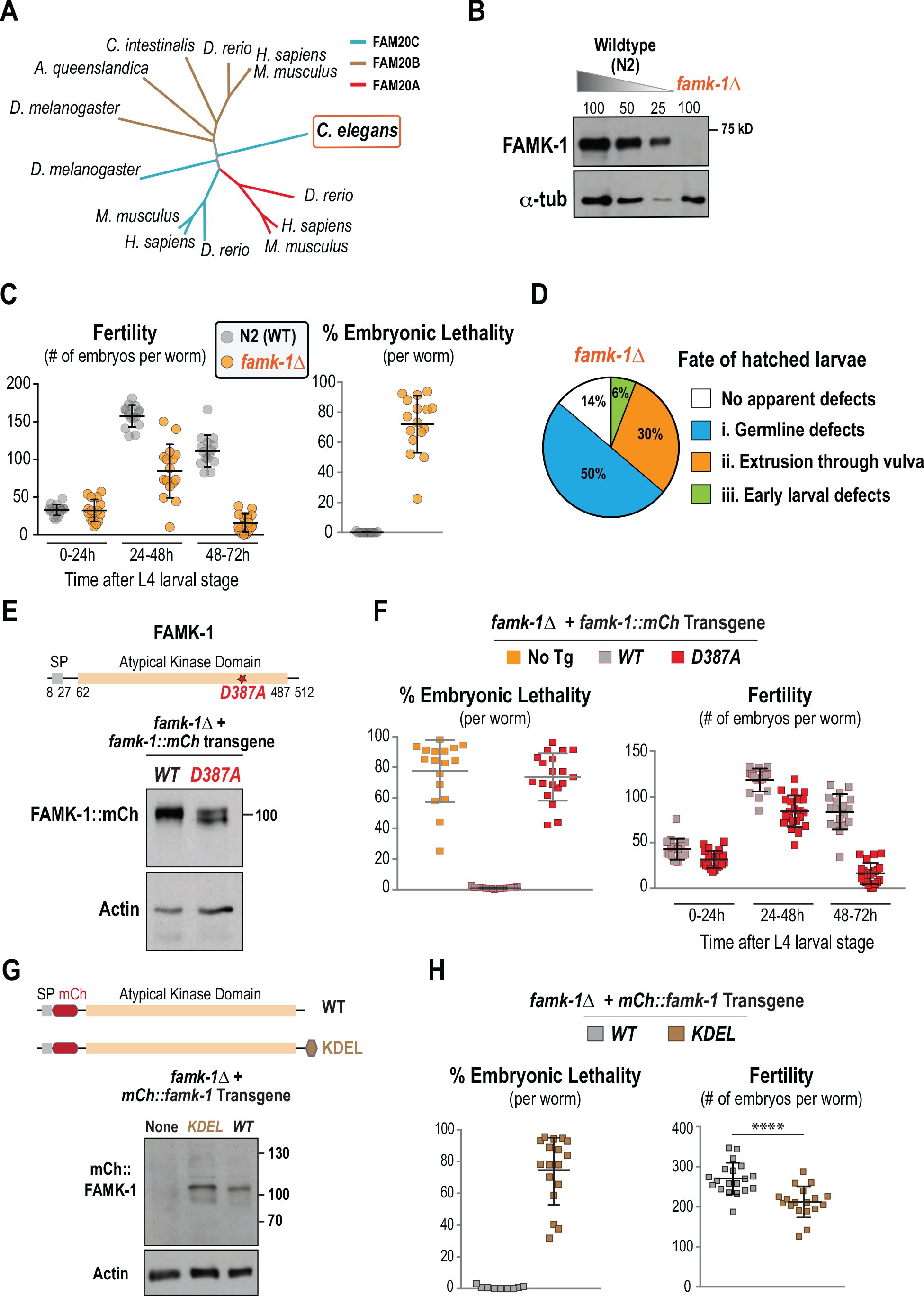
Catalytic activity and presence in the late secretory pathway of FAMK-1 contributes to fertility, embryogenesis and post-hatching development. **(A)** Phylogenetic tree of Fam20 family kinases. Although sequence information alone is unable to assign a specific function to FAMK-1, biochemical analysis has shown it to be a protein kinase. **(B)** I mmunoblot of wildtype (N2) extract and extract prepared from *famk-1∆* worms. atubulin serves as a loading control. **(C)** Fertility and embryonic lethality analysis of individual worms of the indicated genotypes. **(D)** Phenotypes of post-hatching *famk-1∆* larvae. See also *Fig. S1 B*. **(E)** Immunoblot of wildtype and D387A mutant FAMK-1::mCherry expressed from single-copy transgene insertions. The blot was probed using an anti-FAMK-1 antibody. Actin serves as a loading control. **(F)** Fertility and embryonic lethality analysis for the indicated conditions. **(G)** (*top*) Schematics of WT and KDEL mCh::FAMK-1 (with mCherry inserted after signal peptide); (*bottom*) Immunoblot of extracts prepared from the indicated strains probed using an anti-FAMK-1 antibody. Actin serves as a loading control. **(H)** Fertility and embryonic lethality analysis for the indicated conditions.

In humans, mutations in Fam20C cause a spectrum of defects, collectively named Raine Syndrome, that include generalized osteosclerosis, ectopic calcifications, malformed cranium/facial features, and usually result in death shortly after birth (Raine et al. 1989). Patients with less severe Fam20C mutations present with hypophosphatemia, ectopic calcifications and dental anomalies (Fradin et al. 2011; Koob et al. 2011). The role of human Fam20C in the regulation of biomineralization-related processes raises the question as to the role(s) of the Fam20C orthologue in *C. elegans*, an organism with no bones or enamel, and suggests the potential of studying its function in this organism to uncover new roles for this conserved kinase family.

Here, we report the functional characterization of *C. elegans* Fam20C, henceforth referred to as FAMK-1 (for <u>Fam20</u>-like <u>K</u>inase 1). Our results reveal an unexpected role for FAMK-1 in fertility and embryonic development and demonstrate that C-type lectins are substrates of FAMK-1. In addition, the profile of FAMK-1 expression and analysis of its embryogenesis and fertility requirement suggest that FAMK-1 phosphorylation may be important in tissues subject to repeated mechanical strain.

## RESULTS

### The Fam20C Kinase Ortholog FAMK-1 Contributes to Fertility and Development of *C. elegans*

A single *C. elegans* gene, *H03A11.1*, encodes a Fam20-related protein that is analogous to Fam20C as it is a protein kinase with a predicted signal sequence that targets ‘SxE’ motifs for phosphorylation (Xiao et al. 2013). We have named this gene FAMK-1 (for <u>FAM20</u>-like <u>K</u>inase 1). Using CRISPR/Cas9, we engineered a deletion of *famk-1* that removes the majority of the open reading frame (starting from aa 13) and the first 100 bp of the 3’UTR, leading to loss of the gene product **(Fig. 1 B** and **Fig. S1 A**). *famk-1*Δ mutants exhibit reduced fertility and significant embryonic lethality (**Fig. 1 C**). Embryos that hatched into larvae exhibited a diverse spectrum of phenotypes, including morphological defects in early larval stages, multinucleation and compromised germ cell partitions in adult germlines, and extrusion of the intestine through the vulva (**Fig. 1 D** and **Fig. S1 B**). Thus, deletion of *famk-1* is associated with a spectrum of phenotypes, most notably a significant reduction in fertility and embryonic viability.

To confirm that the phenotypes observed were due to loss of *famk-1* function and to test the role of FAMK-1 kinase activity, we replaced endogenous FAMK-1 with transgene-encoded wildtype or kinase-defective versions of FAMK-1. The FAMK-1 crystal structure and biochemical analysis identified D387 (which is homologous to D478 in human *FAM20C*) as being critical for kinase activity (Xiao et al. 2013). We engineered a single copy transgene insertion system to express *famk-1* or *famk-1(D387A)* under control of endogenous *famk-1* promoter and 3’UTR sequences; the transgenes also include the coding region for mCherry fused to the C-terminus (**Fig. 1 E** and **Fig. S1 C**). Immunoblotting verified that FAMK-1::mCherry and FAMK-1(D387A)::mCherry were equally expressed from their respective single copy transgene insertions (**Fig. 1 E**). The wildtype *famk-1::mCherry* transgene fully rescued the embryonic lethality and fertility defects observed in *famk-1*Δ. By contrast, the kinase-defective mutant *famk-1(D387A)::mCherry* exhibited embryonic lethality comparable to *famk-1*Δ (**Fig. 1 F**). The *famk-1(D387A)::mCherry* mutant also exhibited a decline in fertility over time after hatching (**Fig. 1, C and F**) and similar early larval phenotypes as *famk-1*Δ (*not shown*). We conclude that the activity of the secreted kinase FAMK-1 contributes significantly to fertility, embryo viability, and development in *C. elegans*.

### Retaining FAMK-1 in the Endoplasmic Reticulum Phenocopies Loss of FAMK-1

Fam20 kinases have a signal peptide that directs their entry into the secretory pathway. In human cells, Fam20C concentrates in the Golgi and is also secreted into the extracellular environment (Tagliabracci et al. 2012). It is debated whether Fam20 kinases target substrates throughout the secretory pathway and/or extracellularly (Pulvirenti et al. 2008; Tagliabracci et al. 2015). The embryonic viability and fertility phenotypes of *famk-1*Δ motivated us to test whether presence in the secretory pathway was sufficient for FAMK-1 function. For this purpose, we fused the ER retention signal KDEL to the FAMK-1 C-terminus (**Fig. 1 G** and **Fig. S1 D**). To avoid having the tag at the C-terminus, for this analysis we employed transgenes in which mCherry was fused after the signal peptide. The KDEL signal triggers retrograde transport of cargo from the Golgi to the ER, causing proteins bearing it to be retained in the ER (Pulvirenti et al. 2008; Cancino et al. 2014). Immunoblotting confirmed that FAMK-1-KDEL was expressed at comparable levels to wildtype FAMK1 (**Fig. 1 G**). However, in contrast to wildtype FAMK-1, FAMK-1-KDEL failed to rescue the embryonic lethality or fertility defects of *famk-1*Δ (**Fig. 1 H**), and exhibited other defects similar to *famk-1*Δ (*not shown*). In addition, localization analysis indicated that the KDEL sequence altered FAMK-1 distribution and, when expressed in human cells, secretion into the medium of a FAMK-1-KDEL fusion was significantly reduced (*see below*). Thus, while expressed at normal levels, FAMK-1-KDEL does not provide the function of FAMK-1, suggesting that FAMK-1 must be present in the late secretory pathway and/or extracellularly to execute its functions.

### Embryonic Lethality in the Absence of FAMK-1 is Associated with Loss of Inter-Cellular Partitions

We were intrigued by the embryonic lethality observed in *famk-1*Δ, *famk-1(D387A)* and *famk-1-KDEL* as it suggested a new role for secreted protein kinase activity. As a first step, we analyzed the phenotype of *famk-1*Δ embryos that failed to hatch and found that they arrested at multiple stages of embryogenesis, with a significant proportion arresting prior to the morphogenetic events that elongate and shape the embryo into a larva (**Fig. 2 A**). Imaging of *famk-1*Δ embryos expressing a plasma membrane targeting GFP fusion (GFP::PH) and a red fluorescent chromosome marker (mCherry::H2b) revealed that |10% of 16-32 cell-stage and ~45% of 32-64 cell stage *famk-1*Δ embryos exhibited multinucleation (**Fig. 2 B**). In contrast, no multinucleation was observed in over 200 control embryos imaged under similar conditions. Time-lapse imaging of *famk-1*Δ embryos showed that multinucleation was caused by detachment/dissociation of the partitions between adjacent cells (**Fig. 2 C**). 50/62 partition-loss events occurred between sister cells (**Fig. 2 C**; *top row*) 16.2 ± 5.4 min (average ± SD) after full ingression of the cytokinetic furrow. The remainder occurred between cells whose relationship was unclear—they may be products of a cell division event that occurred prior to the start of imaging (**Fig. 2 C**; *bottom row*). Multinucleation is often associated with cytokinesis defects. However, multinucleation was rarely observed prior to the 16-cell stage in *famk-1*Δ embryos (**Fig. 2 B**) and analysis of furrow ingression in one-cell embryos revealed no difference from wildtype (**Fig. S2 A**), suggesting that general cytokinesis mechanisms are not defective in *famk-1*Δ.

**Figure 2.**
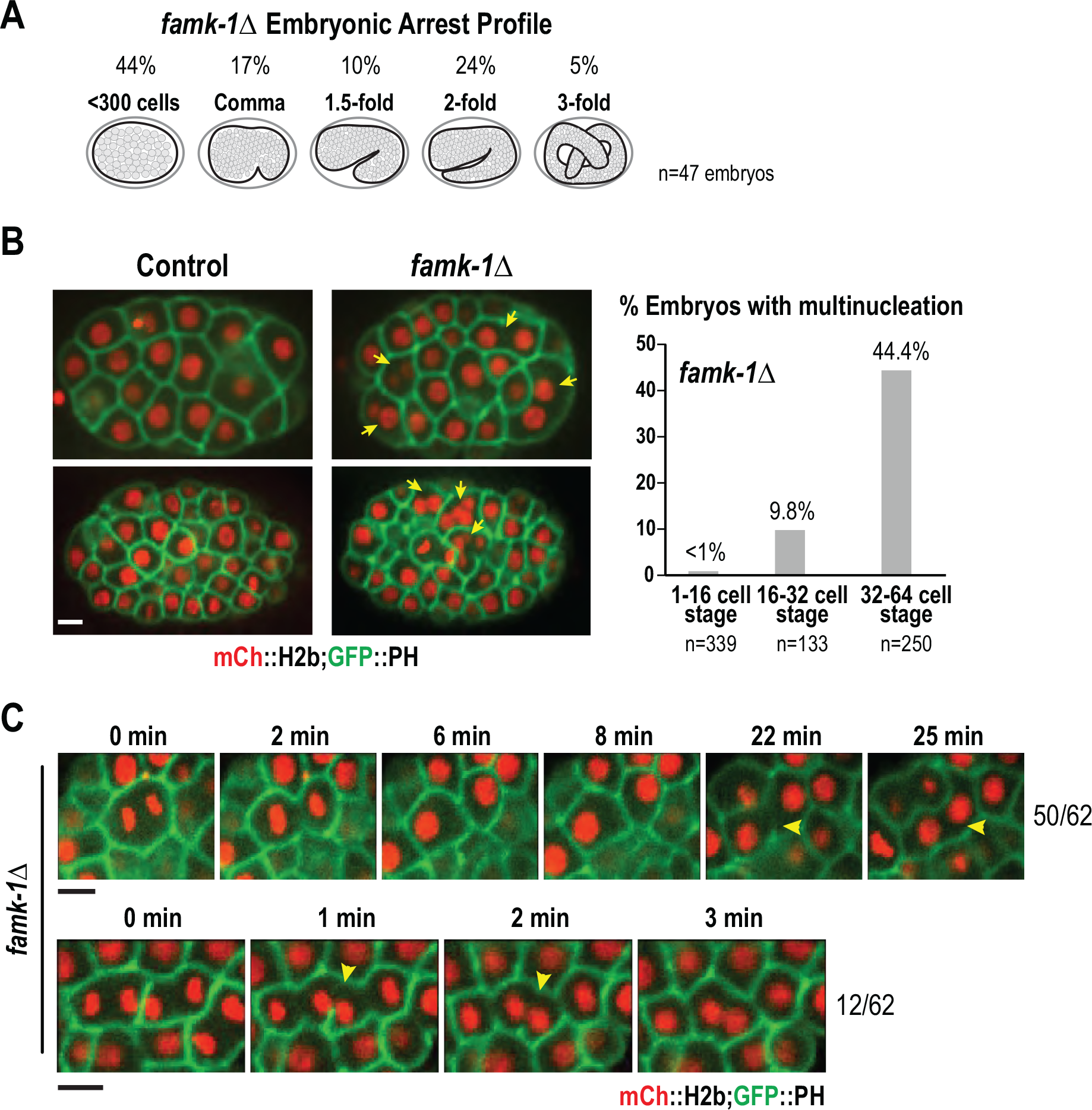
FAMK-1 absence leads to loss of cell partitions and multinucleation in embryos. **(A)** Arrest stage of *famk-1*Δ embryos that failed to hatch. 47 embryos were analyzed. **(B)** Images of control and *famk-1*Δ embryos expressing fluorescent probes that label the plasma membrane and chromatin. Arrows indicated multinucleated cells. Scale bar, 5 µm. The graph on the right shows multinucleation frequency at different stages; to classify an embryo as exhibiting multinucleation, at least 1 clear multinucleation event was required. **(C)** High time-resolution images of *famk-1∆* embryos — the two classes of events associated with multinucleation and their relative frequency are shown. Scale bar, 5 µm.

In addition to the early stages of embryonic development, we also monitored *famk-1*Δ embryos at later stages using live imaging of DLG-1::GFP, which marks epidermal cell boundaries (Koppen et al. 2001). |50% of *famk-1*Δ embryos exhibited epidermal defects, including seam cell defects (mis-positioned or aberrant number, 16/33 embryos) and ventral rupture or enclosure defects (7/33 embryos) (**Fig. S2 B**). These epidermal defects may be a consequence of early multinucleation events or may independently contribute to the observed embryonic lethality. We conclude that there is a high frequency of multinucleation in *famk-1*Δ embryos and that this is followed by significant morphogenesis defects in later stage embryos.

### The Requirement for FAMK-1 During Embryogenesis is Significantly suppressed by Elevated Temperature or Reduced Cortical Tension

The above phenotypic analysis of *famk-1*Δ was conducted at 20°C, the temperature at which *C. elegans* is normally propagated. Temperatures above 25°C cause sterility in *C. elegans* (Byerly et al. 1976; Petrella 2014). A typical approach to enhance mutant phenotypes in genetic models is to induce stress by raising the temperature - in *C. elegans* this is done by placing null mutants at 22-25°C. When conducting such an analysis with *famk-1*Δ embryos, we were surprised to find that elevating the temperature to 23°C significantly suppressed the embryonic lethality of *famk-1*Δ (**Fig. 3 A**); by contrast, fertility was unaffected (**Fig. S3**). Suppression of embryonic lethality at 23°C was also observed with kinase-defective FAMK-1 (**Fig. 3 B**) and ER-retained FAMK-1-KDEL (**Fig. 3 C**). To assess if the temperature increase suppressed the multinucleation phenotype of *famk-1*Δ embryos, we imaged strains with fluorescently marked plasma membrane and chromosomes at 23°C and found that elevated temperature also significantly suppressed multinucleation (**Fig. 3 D**). Thus, the loss of cell partitions and the embryonic lethality in the absence of FAMK-1 is suppressed by increasing the temperature.

**Figure 3.**
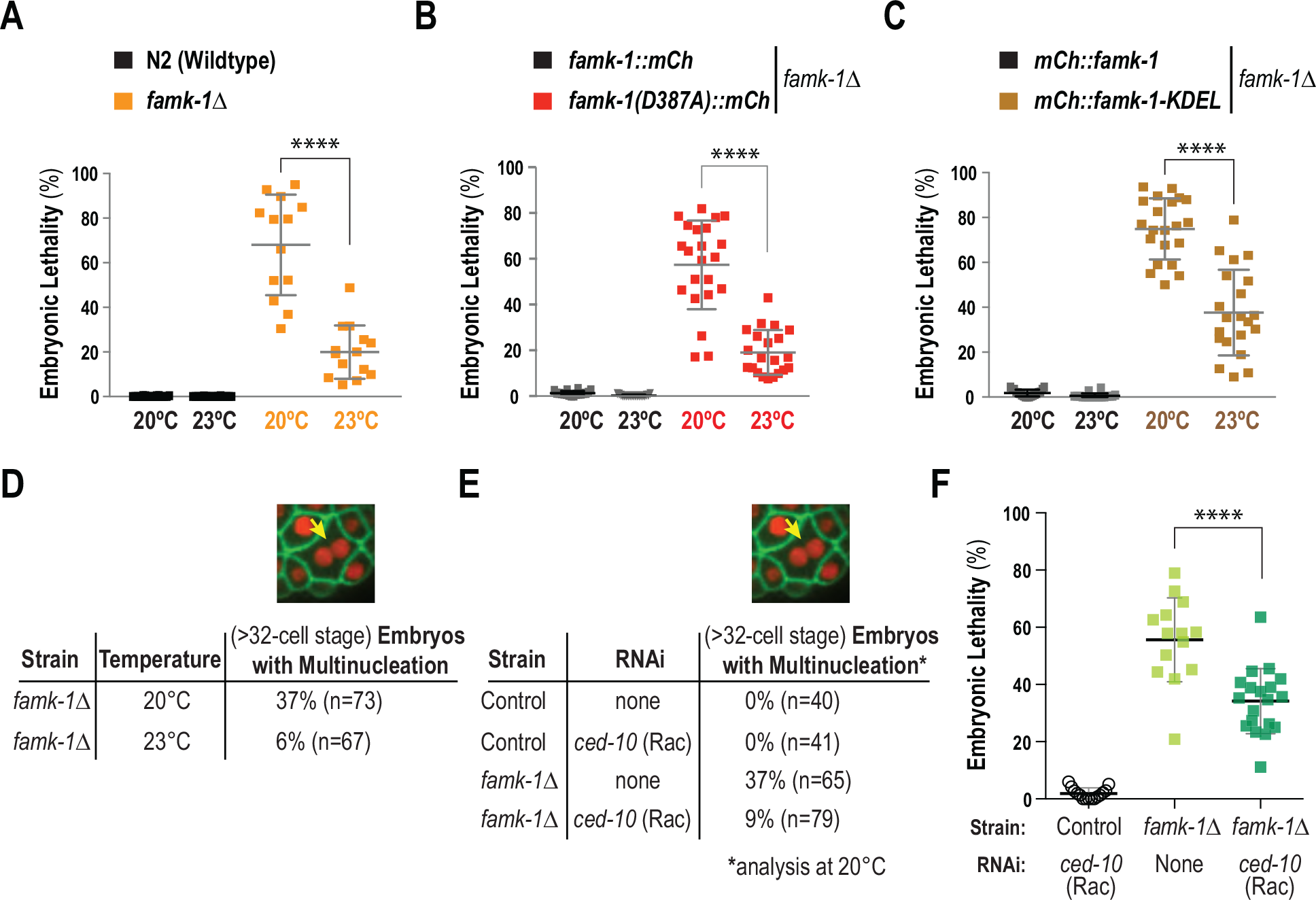
Increased temperature or reduced cortical tension significantly suppress the embryonic lethality observed in the absence of FAMK-1. **(A) – (C) & (F)** Embryonic lethality analysis of the indicated conditions. **(D) & (E)** Analysis of multinucleation in embryos for the indicated conditions. The strains used for the embryonic lethality analysis in *(F)* also expressed GFP::PH and mCherry::H2b markers (the same strains were used for the multinucleation analysis in *(E)*).

The effects of temperature on biological systems are complex, but one possibility that may explain the suppression of multinucleation of *famk-1*Δ, a kinase acting in the late secretory pathway, is that higher temperature makes extracellular interactions between cells more mechanically compliant and less prone to rupture. If this were true, then altering partition stiffness by another means should also suppress the *famk-1*Δ phenotype. To address this possibility, we depleted the major embryonic Rac GTPase, CED-10. Activated Rac directs generation of the Arp2/3-nucleated branched actin network (Pollard 2007). Consequently, Rac inhibition is associated with reduction of effective cortical viscosity and cortical tension (Gerald et al. 1998; Tseng and Wirtz 2004). Similar to elevated temperature, Rac/CED-10 depletion suppressed the multinucleation phenotype (**Fig. 3 E)** and significantly reduced embryonic lethality of *famk-1*Δ (**Fig. 3 F**).

Collectively, the above results suggest that FAMK-1 phosphorylates substrates that likely act extracellularly to ensure that cells placed under mechanical strain in multi-cellular embryos maintain their partitions. Whether loss of partitions is due to regression of incomplete cytokinetic abscission or rupture of abscised but adhered cell-cell boundaries is at present unclear. Notably, the contribution of FAMK-1-mediated phosphorylation in the late secretory pathway to integrity of inter-cellular partitions can be significantly compensated by increased temperature or by reducing the stiffness of the intracellular actin cortex.

### C-type Lectins are Substrates of FAMK-1

To identify FAMK-1 substrates relevant to its function in embryos, we used an informatics approach based on the fact that FAMK-1, like human Fam20C, targets ‘SxE/pS’ motifs (Xiao et al. 2013). Notably, Fam20C progressively phosphorylates a stretch of serines N-terminal to a double glutamate ‘EE’ motif, as phospho-serine is recognized analogously to a glutamate; threonines are not as commonly phosphorylated by this kinase class (Tagliabracci et al. 2015). We therefore searched for gene products with at least three progressive putative FAMK-1 targets (SSSEE). This search yielded 85 hits proteome-wide (**Fig. 4 A**). Refining this list by predicting if the proteins are secreted or anchored by a transmembrane domain on the cell surface narrowed the number down to 43 (**Table S3**). Inspection of this protein list revealed that 13 were lectins: 9 C-type lectins and 4 Galactoside-binding lectins (**Fig. 4 A** and **Fig. S4 A**). Notably, the potential target region in these lectins included more than 3 serines and was followed by a region rich in specific amino acids (H, G, P and R), suggesting shared origin/regulation.

**Figure 4.**
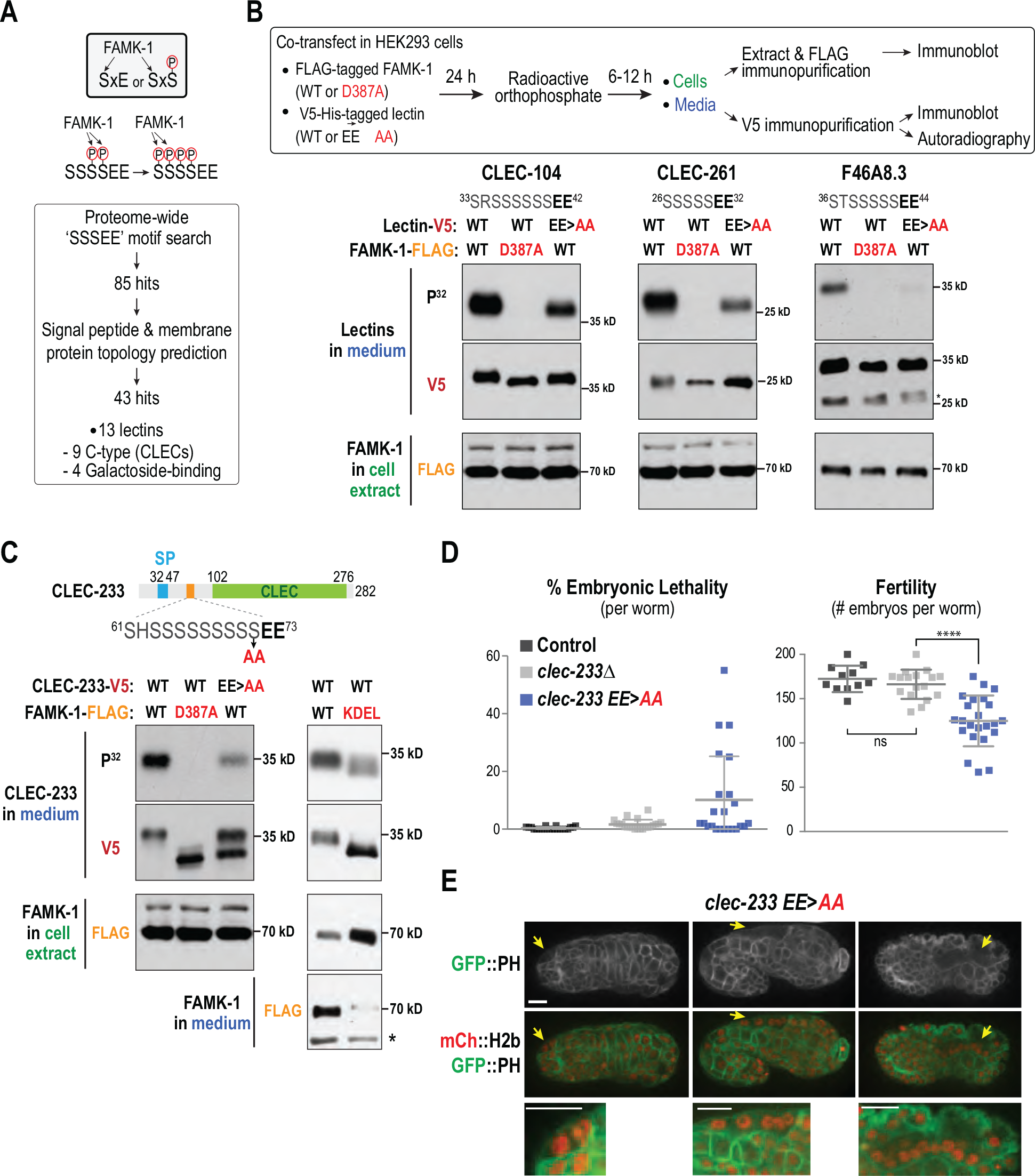
Informatic, biochemical and functional analysis implicates lectins as FAMK-1 substrates. **(A)** Schematics above highlight that the specificity of FAMK-1, which targets serines two residues prior to a glutamate or a phospho-serine, enables progressive phosphorylation of serine clusters preceding two glutamates. The boxed workflow indicates the proteome search approach that revealed lectins as potential FAMK-1 substrates. Signal peptide and membrane topology prediction was performed using SPOCTOPUS and SMART. **(B)** Schematic of experimental protocol employed to test if identified lectins are FAMK-1 substrates. Autoradiograms and blots below show analysis for 3 candidate lectin substrates; the sequence stretches likely to be targets of phosphorylation are shown above the gels. **(C)** Analysis of CLEC-233, conducted as in panel *(B)*. KDEL-containing FAMK-1 was also analyzed for its ability to phosphorylate CLEC-233 (*panels on the right*). **(D)** Embryonic lethality and fertility analysis for the indicated conditions. p<0.0001 is from a t-test. **(E)** Images of late-stage embryos genome-edited to mutate the two glutamates (clec-233 EE>AA) and expressing fluorescent fusions that mark the plasma membrane and chromatin. Arrows point to multinucleation events. Scale bars, 5 µm.

Lectins are proteins that preferentially bind carbohydrates (Mody et al. 1995; Gorelik et al. 2001; Bies et al. 2004; Minko 2004) and are classified into sub-families, including C-type lectins (typically dependent on calcium for carbohydrate binding) and galectins (β-galactoside binding). Both C-type lectins and galectins are found in virtually all organisms. While involved in many biological processes in mammals (Ghazarian et al. 2011), little is known about lectin functions in *C. elegans*. Prior to initiating any genetic analysis, we first tested if FAMK-1 indeed phosphorylated these predicted lectin substrates. We chose 6 of the 13 lectins (5 C-type lectins: CLEC-100, CLEC-104, CLEC-110, CLEC-233, CLEC-261, and 1 galectin: F46A8.3) for this analysis. The coding sequence for each lectin was tagged with a V5-His tag and co-transfected with FLAG-tagged wildtype or D387A kinase-defective FAMK-1 in HEK293 cells. After incubation with radioactive orthophosphate, lectins secreted into the medium were immunopurified and analyzed by immunoblotting and autoradiography; FAMK-1 expression was monitored by preparing cell extracts, immunoprecipitating FAMK-1 and immunoblotting the FLAG tag (**Fig. 4 B**). To specifically test if the serine stretches preceding the two glutamates were sites for phosphorylation, the glutamates were mutated to alanines and the mutant lectins co-transfected with wildtype FAMK-1.

In all six tested lectins, robust phosphorylation was detected in the presence of wildtype but not D387A FAMK-1, indicating that these lectins are phosphorylated by FAMK-1 (**Fig. 4, B and C;** and **Fig. S4 B**). With the exception of CLEC-110, all analyzed lectins were secreted into the medium (**Fig. 4, B and C;** and **Fig. S4 B**). In addition, the EE>AA mutation significantly reduced phosphorylation of all 6 tested lectin substrates; the reduction was evident in the autoradiograms and, for CLEC-100, CLEC-104, and CLEC-233, in the reduced mobility shift relative to the wild-type substrate (**Fig. 4, B and C;** and **Fig. S4 B**). For CLEC-233, we additionally compared wildtype FAMK-1 to FAMK-1-KDEL and observed reduced phosphorylation with the FAMK-1 bearing the ER retention signal; immunoblotting of the medium confirmed that the KDEL sequence reduced secretion of FAMK-1 (**Fig. 4 C**).

Collectively, these results indicate that the lectins identified by the informatic analysis are indeed FAMK-1 substrates and are phosphorylated in the extracellular environment when FAMK-1 is present; in addition, the serine stretches preceding two glutamates identified by our analysis are targets for FAMK-1-dependent phosphorylation.

### CLEC-233 Phosphorylation by FAMK-1 Contributes to the Embryonic Viability and Fertility Functions of FAMK-1

We next wanted to assess if phosphorylation of lectins contributed to FAMK-1 function *in vivo*. For this purpose, we focused on CLEC-233, as inhibition of CLEC-233 was previously reported to lead to mild embryonic lethality in a genome-wide RNAi screen (Kamath et al. 2003). CLEC-233 is predicted to be extra-cellular (**Fig. 4 C**), has an extended FAMK-1 target sequence (^61^SHSSSSSSSSSEE^73^) and an additional ^190^SCE^192^ site. To test if CLEC-233 phosphorylation by FAMK-1 is functionally significant, we generated two mutants: a null allele (*clec-233*Δ) and a genome-edited FAMK-1 phospho-target site mutant, *clec-233(E72A,E73A)*, which significantly reduces phosphorylation by FAMK-1 in the heterologous expression analysis (**Fig. 4 C**). We found that *clec-233(E72A,E73A)*, but not *clec-233*Δ, worms exhibited moderately reduced fertility (**Fig. 4 D**). In addition, a subset of the *clec-233(E72A,E73A)* worms laid a significant proportion of dead embryos (**Fig. 4 D**) – only 1 of 22 *clec-233*Δ worms laid >5% dead embryos; in contrast, 10/23 *clec-233(E72A,E73A)* worms laid >5% dead embryos. Thus, presence of CLEC-233 that is not phosphorylated by FAMK-1 appears more detrimental than the absence of CLEC-233. A potential explanation for this result is that there is redundancy between the large number of C-type lectins in *C. elegans* that compensate for absence of CLEC-233 but not for presence of its poorly phosphorylated form. The embryos laid by only a subset of *clec-233(E72A,E73A)* worms exhibited significant lethality (**Fig. 4 D**), suggesting that CLEC-233(E72A,E73A) induces an organism-wide defect, potentially in the germline. Consistent with this, worms that laid a high proportion of dead embryos also exhibited the most reduced fertility (*not shown*).

To further assess the *clec-233(E72A,E73A)* phenotype, we imaged embryogenesis after crossing in markers to visualize chromosomes and the plasma membrane. |13% of the *clec-233(E72A,E73A)* embryos (7/54) exhibited multinucleation events (**Fig. 4 E**); in a small subset of embryos on plates with high embryonic lethality, there appeared to be near-complete loss of partitions between groups of cells (**Fig. 4 E**). Multinucleation and loss of partitions were observed at much later embryonic stages in *clec-233(E72A,E73A)* than in *famk-1*Δ embryos (compare **Fig. 4 E** to **Fig. 2 B**). Multinucleation and compromised germ cell partitions were also observed in the germline of adult *clec-233(E72A,E73A)* worms (**Fig. S4 C**). These results suggest that CLEC-233 phosphorylation targeting the serine-rich stretch after the signal peptide contributes to fertility and embryogenesis.

### FAMK-1 is Prominently Expressed in the Spermatheca and other Tissues Undergoing Repeated Mechanical Strain

To define the contexts in which FAMK-1 acts in the organism, we generated a transcriptional reporter transgene in which the *famk-1* promoter and 3’UTR control expression of a GFP fusion that concentrates in the nucleus (**Fig. 5 A**). At the L4 larval stage, *famk-1* is transcribed in somatic gonad tissues, prominently in the spermatheca and to a lesser extent in the gonad sheath and vulva, the body-wall muscles and the rectum (**Fig. 5 A**). *famk-1* expression was also detected in oocytes in the adult gonad (**Fig. 5 B**). Although transcriptional reporter signal was not detected in embryonic nuclei (*not shown*), FAMK-1 protein was detected by immunoblotting of embryo extracts (**Fig. 5 C**), suggesting that maternally loaded protein/mRNA contributes to the embryonic functions of FAMK-1 analyzed above.

**Figure 5.**
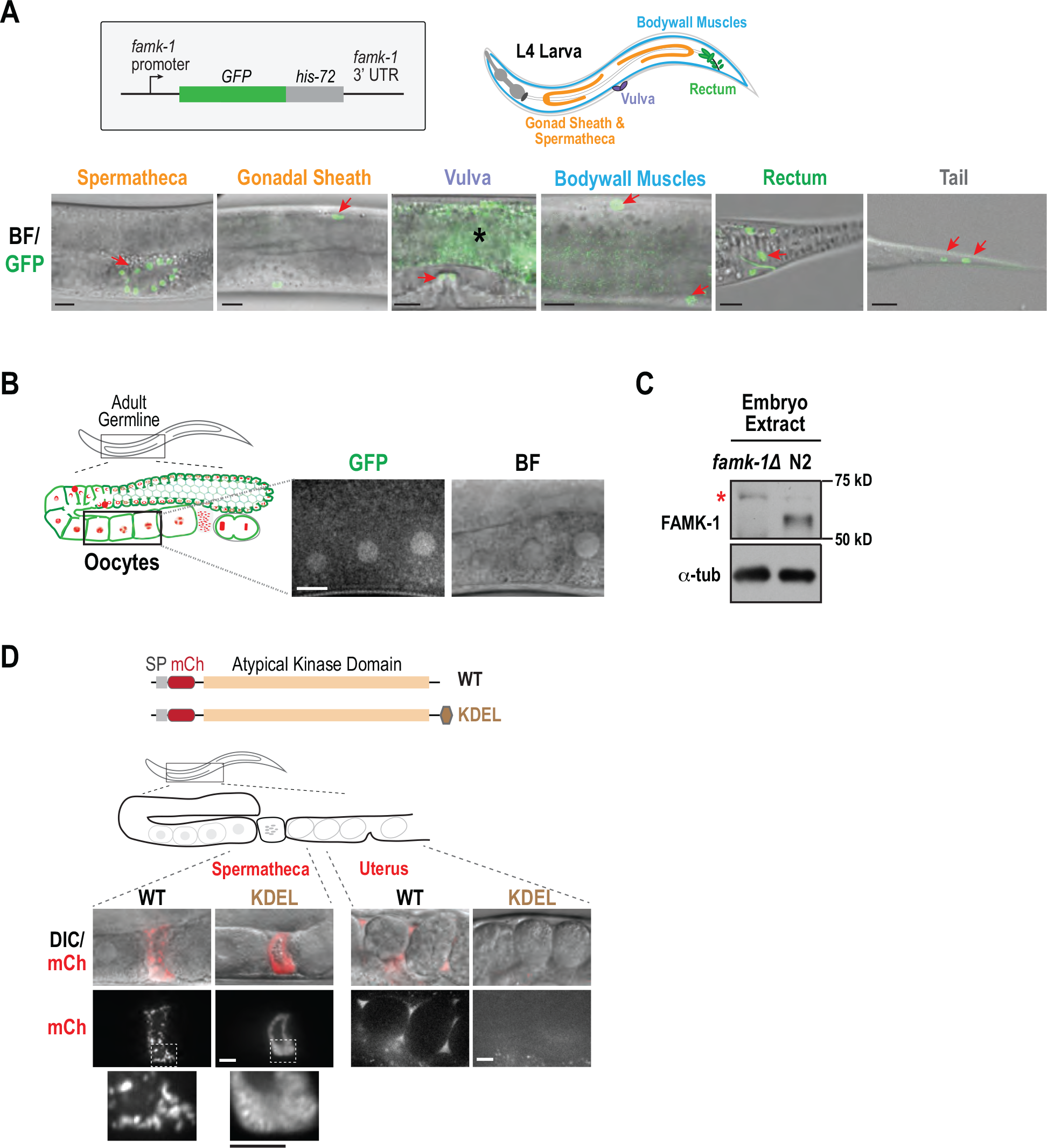
Pattern of FAMK-1 expression assessed using transcriptional and translational reporters. **(A)** Schematic shows transcriptional reporter used to analyze tissue distribution of FAMK-1 in the L4 larval stage. The schematic on the right highlights the tissues where FAMK-1 expression was observed. Images below show overlays of brightfield (BF) and nuclear GFP signals; arrow highlight fluorescent nuclei. The most prominent expression of FAMK-1 in the L4 larva (and adult) is in the spermatheca. Asterisk marks background autofluorescence. Scale bar, 10 µm. **(B)** Image of oocytes in an adult highlighting FAMK-1 expression in the germline. Scale bar, 10 µm. **(C)** Immunoblot of embryos purified from the indicated strains. Asterisk marks a background band. a-tubulin serves as a loading control. **(D)** Comparison of mCherry-fused WT and KDEL FAMK-1 in the spermatheca and the uterus. For images in other tissues, see *Fig. S5*. Scale bar, 10 µm.

In addition to the transcriptional reporter, we also monitored localization of mCherry fusions of wildtype FAMK-1 and FAMK-1-KDEL. This analysis confirmed FAMK-1 expression in tissues highlighted by the transcription reporter (**Fig. S5**). Notably, the pattern of FAMK-1 localization was consistently different between wildtype FAMK-1 and FAMK-1-KDEL (**Fig. 5 D** and **Fig. S5**). In the spermatheca and the vulva, wildtype FAMK-1 appeared patchy throughout the organ whereas FAMK-1-KDEL was localized in a mesh-like pattern around nuclei (**Fig. 5 D** and **Fig. S5**). In the uterus, wildtype FAMK-1 was present in the luminal space, but no FAMK-1-KDEL was observed (**Fig. 5 D**), consistent with its not being secreted. Interestingly, the KDEL fusion revealed FAMK-1 expression in two locations not detected with the transcriptional reporter: the head and a specific cell adjacent to the vulva (potentially the CANL or CANR neuron) (**Fig. S5**). One reason why these expression sites were missed in the transcriptional reporter analysis is that the level of transcription was too low to generate nuclear GFP signal detectable over background autofluorescence.

Taken together, the transcriptional reporter and protein localization analysis indicates FAMK-1 expression in multiple tissues in the adult, which likely underlie the post-hatching defects observed in *famk-1*Δ worms. Notably, many of these tissues, e.g. the spermatheca and the vulva, are subject to repeated mechanical strain as are cells in developing embryos where FAMK-1 helps maintain cellular partitions. This pattern of expression suggests potential FAMK-1 phosphorylation of extracellular substrates may be important to withstand this strain.

### Selective Expression of FAMK-1 in the Spermatheca Rescues the Fertility Defect of *famk-1*∆

The most prominent localization observed for FAMK-1 in the adult is in the spermatheca, the organ responsible for ovulation and fertilization in *C. elegans* hermaphrodites. Mating with wildtype males did not rescue the fertility defect of *famk-1*Δ (**Fig. S6 A**), suggesting that compromised spermathecal function may contribute to this phenotype. To test this possibility, we wanted to express FAMK-1 selectively in the spermatheca in the absence of endogenous FAMK-1 and assess the consequences on fertility and embryo viability. For this purpose, we decided to use the *sth-1* promoter (Bando et al. 2005) and the *famk-1* 3’UTR; expression of GFP::HIS-72 (histone H3.3) under control of these regulatory elements from a single copy transgene insertion was observed prominently in spermathecal nuclei and two vulval nuclei, and not in any other tissues in the L4 larva/adult (**Fig. 6 A**). Immuoblotting of *famk-1*Δ worms expression p*famk-1* versus p*sth-1*-driven *famk-1* confirmed FAMK-1 expression under control of p*sth-1* (**Fig. 6 B**); as expected given the broad distribution of endogenous FAMK-1, the expression level of p*sth-1*-driven FAMK-1 was lower. *famk-1* expressed under control of the *sth-1* promoter rescued the fertility defect observed in *famk-1*Δ (**Fig. 6 C**) and reduced the embryonic lethality of *famk-1*Δ (**Fig. 6 C**). The extrusion through vulva phenotype, indicative of loss of vulval integrity (Reinhart et al. 2000; Leiser et al. 2011; Leiser et al. 2016), was also rescued by p*sth-1*-driven FAMK-1 (*not shown*), potentially due to the *sth-1* promoter being active in specific vulval cells (**Fig. 6 A**). By contrast, morphological defects at early larval stages were observed at a similar frequency to *famk-1*Δ (*not shown*). Overexpression of p*sth*-1-driven FAMK-1 from extrachromosomal arrays also restored brood size but embryonic lethality was only mildly reduced (**Fig. S6 B**). Thus, spermathecal expression of FAMK-1 supports the fertility function of *famk-1*Δ. We note that FAMK-1 is secreted and the eggshell surrounding the embryo forms after fertilization, which occurs during passage through the spermatheca. Thus, some spermathecally expressed and secreted FAMK-1 may end up in the extracellular space between the embryo plasma membrane and the outer eggshell layers, which could contribute to the reduction in embryonic lethality.

**Figure 6.**
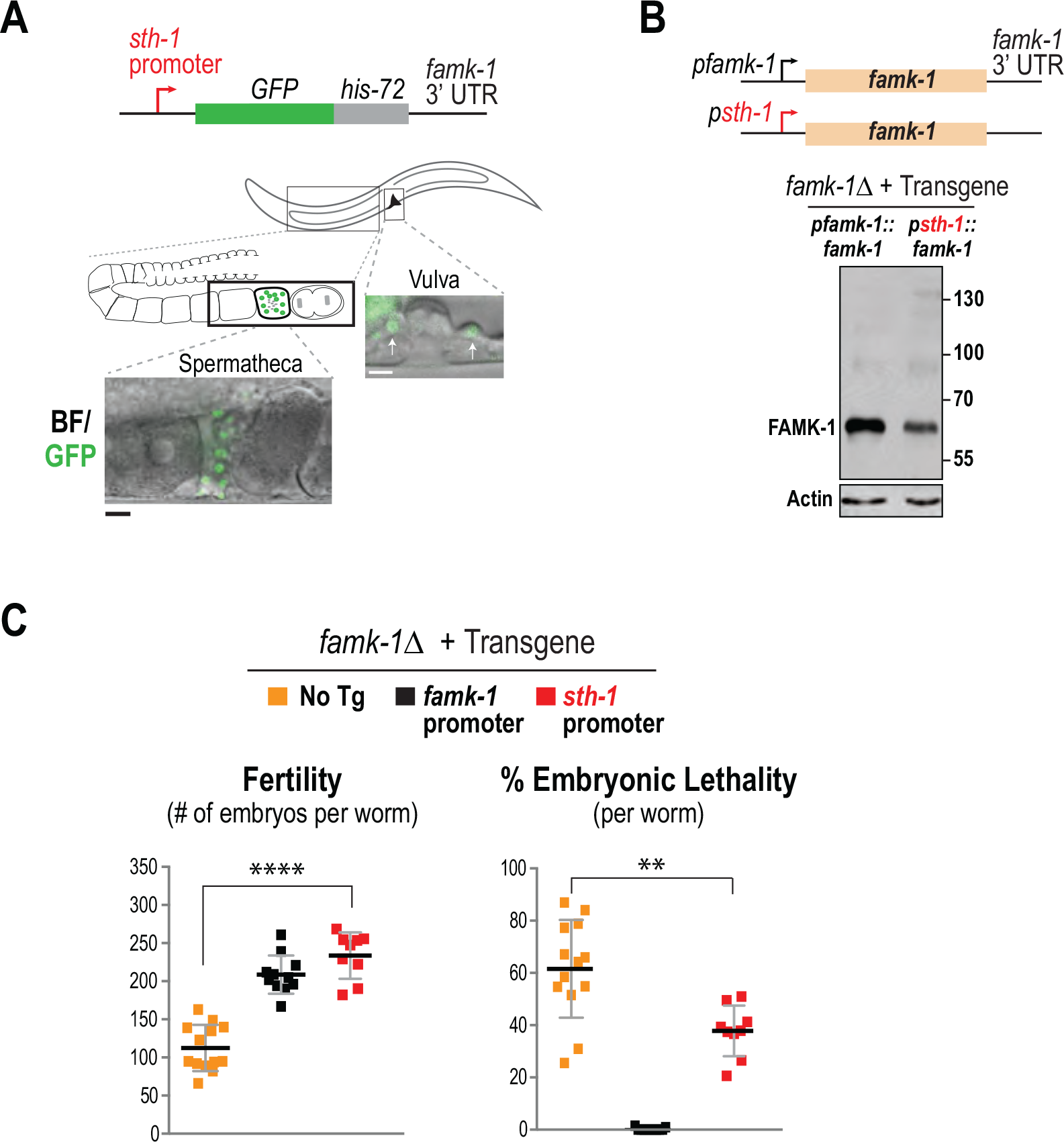
Selective expression of FAMK-1 in the spermatheca restores normal fertility to *famk-1*Δ worms. **(A)** Images of a nuclear GFP reporter expressed under control of the spermathecal promoter p*sth-1*. The reporter shows strong expression in the spermatheca and weaker signal in 2 cells in the vulval region; no additional expression was observed. Scale bars, 10 µm. **(B)** Immunoblot of the products of the two transgenes schematized above in the absence of endogenous FAMK-1. Actin serves as a loading control. **(C)** Embryonic lethality and fertility analysis forthe indicated conditions. ****p<0.0001 and **p<0.01 from t-tests.

The spermatheca is a contractile tube composed of 24 smooth muscle-like cells, separated from the gonad arm by a constriction and from the uterus by a valve (Kimble and Hirsh 1979; Ward and Carrel 1979). The spermatheca plays a critical physical role in ovulation, by accommodating the ovulating oocytes during fertilization and pushing the fertilized oocytes into the uterus while constricting. Spermatheca constriction depends on calcium pulses, which occur during the ovulation cycle (Kovacevic et al. 2013). During the first few ovulations, no reproducible difference in calcium pulses was observed between wildtype and *famk-1*Δ worms (**Fig. S6 C**), indicating that there is no significant defect in spermathecal structure or inter-cellular signaling in the absence of FAMK-1. In support of this, the fertility defect of *famk-1*Δ worms is gradual (**Fig. 1, C and F**). We conclude that FAMK-1 expression in the spermatheca contributes to fertility, potentially by supporting this mechanically highly active structure. The reduced fertility of *clec-233(E72A,E73A)* additionally suggests that CLEC-233 phosphorylation by FAMK-1 contributes to spermathecal function.

## DISCUSSION

### New Functions for the Secreted Protein Kinase Family

The field of secreted kinases originated with the finding that *FAM20C*, the gene mutated in Raine Syndrome, encoded a kinase that was sufficiently distinct in primary sequence from canonical kinases to have been left off the kinome tree (Manning et al. 2002; Johnson and Hunter 2005). Mammalian Fam20C phosphorylates secreted proteins at SxE/pS motifs, notably calcium-binding proteins such as casein, osteopontin, and members of the secretory calcium-binding phosphoprotein (SCPP) family (Ishikawa et al. 2012; Tagliabracci et al. 2012; Lindberg et al. 2015). While Fam20C is ubiquitously expressed, it is present at higher levels in mineralized tissues and mutations in humans result in Raine syndrome, an extremely rare, autosomal recessive osteosclerotic bone dysplasia (Raine et al. 1989; Simpson et al. 2007). Ablation of Fam20C expression in mice results in bone lesions and dental abnormalities among other phenotypes (Wang et al. 2010; Vogel et al. 2012). Taken together, the most notable functions of Fam20C in higher eukaryotes involve calcium-binding proteins in mineralized tissues. However, inspection of the secreted phosphoproteomes of cerebrospinal fluid, human serum and plasma reveals that the majority of their phosphoproteins contain phosphate within SxE/pS motifs and that these phosphoproteins have no apparent roles in biomineralization (Bahl et al. 2008; Zhou et al. 2009; Carrascal et al. 2010; Salvi et al. 2010). Furthermore, depletion of Fam20C in a variety of cell types uncovered more than 100 secreted phosphoproteins as genuine Fam20C substrates whose annotations span a broad range of biological processes (Tagliabracci et al. 2015). Thus, Fam20C is likely to act broadly in phosphorylation of extracellular proteins and contribute to as-yet-unknown functions.

In order to address roles of Fam20C not involving biomineralization, we initiated a study of Fam20C in *C. elegans*, an organism that shares a common ancestor with humans |500-600 million years ago and has no bones or teeth. Notably *C. elegans* has one Fam20C-like predicted protein, FAMK-1, which has been structurally and biochemically characterized to be a protein kinase targeting SxE/pS motifs (Xiao et al. 2013). Our analysis indicates that FAMK-1 has prominent functions in embryogenesis and fertility in *C. elegans* and that lectins are a substrate class that contributes to these functions. We also demonstrate the importance of FAMK-1 transit into the late secretory pathway for its function and document expression of FAMK-1 in multiple tissues as well as phenotypes observed in larval and adult stages. Collectively, our findings suggest that phosphorylation of extracellular substrates by FAMK-1 may be important in contexts where tissues undergo repeated mechanical strain. In light of this proposal, it is interesting to note that a link between Fam20C phosphorylation and prevention of cardiac arrhythmia has been recently reported in humans (Pollak et al. 2017).

### FAMK-1 Must Transit into the Late Secretory Pathway to Execute its Functions

The generation of a *famk-1* knockout mutant in *C. elegans*, the definition of a phenotypic profile for the knockout, and the ability to employ single copy transgene insertions to replace endogenous FAMK-1 with any engineered version has enabled us to address an important question in the secreted kinase field—where in the secretory pathway is phosphorylation of functionally critical substrates occurring? In human cells, fusion of the ER retention signal KDEL to Fam20C did not prevent phosphorylation of selected secreted substrates in tissue culture cells, suggesting that substrate phosphorylation can occur in the lumen of the ER (Tagliabracci et al. 2015). However, whether this extent of phosphorylation was sufficient for Fam20C function has remained unclear.

Using our replacement system to express FAMK-1 fused to KDEL as the sole source of FAMK-1, we found that the ER-retained FAMK-1, while expressed at normal levels, exhibited nearly identical phenotypes as the *famk-1*-null mutant and a kinase-defective FAMK-1 mutant. This result suggests that FAMK-1 activity is critical in the late secretory pathway, specifically the Golgi, and possibly also in the extracellular environment. Unlike the recently identified secreted kinases, the existence of multiple secretory pathway phosphatases has been known for decades (Gutman 1959; Kiefer 1977; Goldfischer 1982; Ek-Rylander et al. 1994; Bull et al. 2002; Rigden 2008). Thus, we postulate that FAMK-1 substrates in the Golgi and outside the cell are regulated by phosphorylation/ dephosphorylation, in a manner similar to cytoplasmic proteins, and that restriction of FAMK-1 to the ER shifts the balance to dephosphorylation, thereby phenocopying loss of FAMK-1 activity.

### FAMK-1 Expression and Phenotypic Spectrum: A Role for Extracellular Phosphorylation by FAMK-1 in the Function of Mechanically Strained Tissues?

FAMK-1 is expressed in multiple tissues and has a complex phenotypic spectrum, which is not surprising given that the loss of FAMK-1 catalytic activity likely affects phosphorylation of a potentially large number of extracellular substrates in different tissues. Based on the pattern of expression and the phenotypes observed, we propose that one important role of FAMK-1 phosphorylation is to maintain functionality of tissues/organs under repeated mechanical strain. The most prominent expression of FAMK-1 is in the spermatheca, an organ that undergoes repeated cycles of extension and relaxation during ovulation (one ovulation event occurs every 20 minutes in young adult hermaphrodites). While spermathecal structure and early function was normal in the absence of FAMK-1, embryo production declined rapidly over time, suggesting an exhaustion-type defect. A second example is the |30% of adult *famk-1*Δ worms that exhibit extrusion of the intestine through the vulva; the vulva is another organ that undergoes repeated cycles of mechanical strain during egg-laying. Notably, the *famk-1* transgene with the *sth-1* promoter, selected to drive expression in the spermatheca, was also expressed in vulval cells and not only rescued the fertility defect of *famk-1*Δ but also rescued the extrusion through vulva phenotype. These observations suggested that FAMK-1 activity is important in the context of repeated mechanical strain.

In embryos lacking FAMK-1 activity, the major defect observed was multinucleation; this phenotype was observed at stages when cells are likely under mechanical strain from neighbors. Notably, the multinucleation (and the associated embryonic lethality) observed in the absence of FAMK-1 activity was significantly reduced by either temperature elevation or by reduction of intracellular cortical tension (by depletion of the actin-regulating GTPase Rac). These results lend support to the idea that FAMK-1 phosphorylation of extracellular substrates is important in mechanically strained contexts. Precisely how FAMK-1 phosphorylation acts in these contexts will require future work to elucidate. We note that temperature elevation did not suppress the fertility defect of *famk-1*Δ and, while FAMK-1 is expressed in body wall muscles, we have not observed obvious defects in movement of the fraction of worms that lack other phenotypes. We have also not investigated FAMK-1 function at other sites where it is expressed. Thus, in addition to testing the hypothesis suggested by our analysis of FAMK-1 function in the spermatheca, the vulva, and during embryogenesis, it will be important to conduct functional assays for other tissues/cell types in which FAMK-1 is expressed to understand its functions at those sites.

### Lectins are Functionally Relevant Substrates of FAMK-1

An important question raised by our analysis in embryos is how a secretory pathway kinase functions to prevent loss of inter-cellular boundaries. One potential explanation is that FAMK-1 phosphorylates adhesion molecules that reside along cell-cell contacts to alter their properties. In human cells, adhesion molecules, including cadherin-2 (CDH-2) and amyloid beta A4 protein (APP), are substrates of Fam20C (Tagliabracci et al. 2015). In *Drosophila*, Four-Jointed (Fj), the first Fam20-like kinase to be identified, phosphorylates the Cadherin domains of Fat and Dachsous adhesion molecules (Ishikawa et al. 2008). However, while depletion of the major *C. elegans* epithelial-cadherin (HMR-1) causes epidermal enclosure defects and embryonic arrest (Costa et al. 1998; Raich et al. 1999), it does not result in multinucleation similar to what is observed with loss of FAMK-1 (*not shown*). Analysis of other *C. elegans* cadherins is more limited (Pettitt 2005), but they have not been reported to contribute to embryo viability.

To identify FAMK-1 substrate(s) in an unbiased manner, we used the fact that FAMK-1 progressively phosphorylates serines preceding two glutamates to identify potential substrates. This search identified lectins, proteins with carbohydrate-recognition domains, as potential FAMK-1 substrates. Lectin function in *C. elegans* is poorly studied (in part because of the very large expansion of lectins in this organism – there are over 200 lectin-encoding genes (Takeuchi et al. 2008) – but in mammals, lectins are known to be involved in cell-cell contacts, lipid binding, and the immune response (Ghazarian et al. 2011). We focused on the lectins because we reasoned that the interaction of these proteins either with other proteins or carbohydrate moieties on extracellular matrix proteins could influence mechanical properties. Using heterologous expression in human cells, we confirmed that all six tested lectins were FAMK-1 substrates and that mutation of the double-glutamate site reduced phosphorylation.

Functional analysis indicated that elimination of FAMK-1 phosphorylation sites in one lectin (CLEC-233) resulted in phenotypes observed when FAMK-1 function is compromised, including multinucleation in embryos, multinucleation in adult germlines and embryonic lethality. Lectins are increasingly recognized as critical players in diverse aspects of multicellular life (Brown et al. 2018). In particular, there is growing evidence for potential functions for C-type lectins in bone mineralization and/or mineralized structure formation (Neame et al. 1992; Wewer et al. 1994; Aspberg et al. 1999; Mann and Siedler 2006; Mann et al. 2008; Mann et al. 2010; Yue et al. 2016; Flores and Livingston 2017). Specifically, C-type lectins of the mammalian OCIL family inhibit osteoclast formation (Zhou et al. 2002; Nakamura et al. 2007; Kartsogiannis et al. 2008). OCIL members resemble *C. elegans* CLEC-233 in that they are composed of an intracellular domain followed by a transmembrane domain and an extracellular C-type lectin-like domain that contains an “SxE” motif (Zhou et al. 2002), making them potential Fam20C substrates. Future work will be needed to test if this class of C-type lectins is indeed phosphorylated by Fam20C and whether this phosphoregulation contributes to the functions of Fam20C in mammals.

In summary, we present here functional analysis of the Fam20C-related secreted kinase FAMK-1 in *C. elegans*, an organism that lacks the structures (bones and teeth) most prominently associated with the function of Fam20C in mammals. Our findings highlight the multiple different contexts in which FAMK-1 acts and demonstrate that it must transit into the late secretory pathway to execute its functions. In addition, we have discovered a new family of secreted substrates for FAMK-1, the lectins. The tools we established should enable future systematic analysis of target substrates and potential functions of their extracellular phosphorylation in all of the different contexts where this secreted kinase is produced.

## ACKNOWLEDGMENTS

We thank members of the Dixon, Desai and Oegema labs for helpful discussions and Coleman Clifford for technical support. This work was supported by grants from the NIH (DK018849-41 & DK018024043 to J.E.D. and C.A.W; GM074215 to A.D.). A.G-G. acknowledges support from an EMBO Long Term Fellowship (ALTF 251-2012) during the initial stages of this work. A.D. & K.O. acknowledge salary and other support from the Ludwig Institute for Cancer Research.

## MATERIALS AND METHODS

All strains used in the study are listed in **Table S1**.

### Genome-edited strains

The *famk-1*Δ allele (*famk-1(lt33::cb-unc-119+)*, OD2102) was generated using CRISPR/ Cas9. A repair plasmid pOD2744/pAG58 was used, which contains the *Cb-unc-119* selection marker and *famk-1* locus homology arms. The 5’ homology arm is 3067 bp long, composed of 3041 bp upstream of the *famk-1* ORF and the first 26 bp of the ORF. The 3’ homology arm is 1687 bp, beginning 190 bp downstream of the *famk-1* stop codon. A mix of two guides was used:

5’*cgattattcactttggcgat*3’ (N-terminus) and 5’*tggatgggataccttcttg*3’ (3’UTR). The knockout strain was genotyped and backcrossed 7 times to N2 (wildtype).

A co-CRISPR strategy (Arribere et al. 2014; Kim et al. 2014) employing the *dpy-10* marker was used to generate the *clec-233* null mutant and the *clec-23(EE>AA*) (*clec-233 E72A,E73A*) mutant allele. A 1:4 mix of *dpy-10*:*clec-233* crRNA (IDT,100 µM final) was prepared. The *dpy-10* target sequence was 5’*gctaccataggcaccacgag*3’. For *clec-233* deletion, a mix of two target sequences was used: 1-5’*atgtacgtatgtggaaagag*3’ 2-5’*gtttttttacaatcccggtt*3’. For *clec-233 EE>AA*, the target sequence was 5’*gctcatcgtccgaagagca*3’. The crRNA:trRNA duplex was generated by 5 minutes incubation at 95°C followed by 5 minutes incubation at room temperature. A mix of 2.2 µl of crRNA:trRNA duplex, 4.8 µl of 40 µM purified *S. pyogenes* Cas9-NLS protein (a gift from K. Corbett) and 1 µl (1.2 µg) of ssDNA repair template (Valuegene Inc).

<u>For *clec-233*Δ, sequence of repair template:</u>

5’***ggtatactctactaacttagacaattacaacaaaaaaatacgaaagtttgaataatttattggcatgcttaacgt**TTAA CGTGATACACCCCTC<u>accgggattgtaaaaaaacagaaaattgtggttaac</u>*3’.

Bold– sequence downstream of *clec-233* open reading frame (in 3’UTR); Capitalized– sequence at 3’ end of *clec-233* open reading frame; Underlined– sequence ending 7 bp upstream of *famk-1* open reading frame (in 5’UTR region).

<u>For *clec-233(EE>AA)*, sequence of repair template:</u>

5’*gtctatgtggtctatgtggtggtctatggggcctgtgtggcggtctatgatggccatg<u>cgctgc</u>ggacgatgagctagacgaag acgaagagtgggagtaatagcctcccccacctccaccacct*3’ (underlined nucleotides introduce the E72A and E73A mutations)

The final mix was injected into gonads of OD95 gravid adult worms (which carry transgenes expressing GFP::PH(PLC1delta1) and mCherry::HIS-58 markers). Injected worms were singled on plates and maintained at 23°C. Three days later, F1 progeny were screened and ~40 Roller worms were singled on plates, allowed to lay eggs, and then screened by PCR for presence of *clec-233* deletion or introduction of the point mutations.

For genotyping the *clec-233* deletion, a combination of three oligos was used: Forward – 5’*ccatgtcaatgttctgagac*3’ (5’UTR); Reverse 1-5’*ggcgcagaaactcttgctt*3’ (ORF); Reverse 2-5’*gagtgaggaagttgtacatg*3’ (3’UTR). When *clec-233* is deleted, the 5’UTR and 3’UTR primers generate a 696 bp PCR product, while when there is no deletion, the 5’UTR and ORF primers generate a 484 bp product.

For genotyping *clec-233(EE>AA)*, the following oligos were used for PCR screening: Forward-5’*agctcatcgtccgcagcg*3’ Reverse-5’*aggttatcaggttcgtcacg*3’. Presence of mutations E72A, E73A yields a 503 bp band, while lack of mutations yields no band. Mutation insertion was further verified by amplifying the genomic region between oligos 1–5’ccatgtcaatgttctgagac3’ and 2–5’aggttatcaggttcgtcacg3’, followed by sequencing (Retrogen Inc) with oligo 5’*aagcaagagtttctgcgcc*3’.

### Single copy transgene insertions of famk-1

Single copy transgene insertions were generated using a transposon-based strategy (MosSCI:, (Frokjaer-Jensen et al. 2008)). All *famk-1* transgenes included 2701 bp upstream of the *famk-1* start codon (*famk-1* promoter) and 1761 bp downstream of the *famk-1* stop codon (*famk-1* 3’UTR). Both untagged and mCherry/GFP-tagged transgenes were constructed. The different constructs were cloned into vectors with the *Cb-unc-119* selection marker: pCFJ151 for ChrII insertion (*ttTi5605*) or pCFJ352 for ChrI insertion (*ttTi4348*), then injected into strains EG6429 (*ttTi5605*, Chr II) or EG6701 (*ttTi4348*, Chr I) in a mix together with a Mos1 transposase coding plasmid (pJL43.1, P*glh-2::Mos1 transposase*, 50 ng/µl) and three plasmids encoding fluorescent markers for negative selection (pCFJ90 [P*myo-2::mCherry*, 2.5 ng/µl], pCFJ104 [P*myo-3::mCherry*, 5 ng/µl] and pGH8 [P*rab-3::mCherry*, 10 ng/µl]). After 7-10 days, moving progeny with no fluorescent markers were selected and transgene insertion was confirmed by PCR.

For comparison of wildtype and D387A mutant of *famk-1*, the coding sequence for mCherry (optimized for expression in *C. elegans*) was inserted immediately before the stop codon.

For N-terminal tagging of *famk-1*, the mCherry sequence was introduced immediately after the signal peptide (i.e. after aa 27) using a synthesized G-block DNA fragment (Integrated DNA Technologies, IDT). Two DNA fragments were synthesized then annealed to produce the desired fragment (the sequence was broken into two fragments to break repeats which complicate G-block gene synthesis):

Fragment 1:

5’(tgcatatctccggtgcgtaacatgcagtac)tggagtaaattattaattaatcgctttgatctcttgtttcctcattctaccccattattt atctataatagcttatgaacctcaccccataaatgctcattttgcagtgtcacaaaactgggagccggcgtttgatgaagctgaaa ccagcaaagagcaagaagtgagacgagaaaagtccagaaagac*atgcggtgcaatataaagcgattattcactttggcgat cggagtttttgcggctacactggttataatctcgttttccaag***ggtggtggtggtggaggt**tctgtctcaaagggtgaagaagata acatggcaattattaaagagtttatgcgtttcaaggtgcatatggagggatctgtcaatgggcatgagtttgaaattgaaggtgaa ggagaaggccgaccatatgagggaacacaaaccgcaaaactaaaggtaagtttaaacatatatatactaactaaccctgatt atttaaattttcaggtaactaaaggc<u>ggaccattaccattcgcctgggacatcctct</u>3’

Fragment 2:

5’<u>gaccattaccattcgcctgggacatcctct</u>ctccacagttcatgtatggaagtaaagcttatgttaaacatccggcagatatacc agattatttgaaactttcattcccggagggttttaagtgggaacgcgtaatgaattttgaagacggaggagttgttacagtgacgca agactcaaggtaagtttaaacagttcggtactaactaaccatacatatttaaattttcagcctccaagatggagaatttatttataaa gtcaaacttcgaggaacgaatttcccctcggatggacctgttatgcagaagaagactatgggatgggaagcttcaagtgaaag aatgtaccctgaagacggtgctcttaagggagagattaaacaacgtcttaaattgaaagatggaggacattacgatgctgaggt aagtttaaacatgattttactaactaactaatctgatttaaattttcaggtgaagacaacttacaaagccaaaaaaccagttcagct gccaggagcgtacaatgttaatattaaactggatatcacctcccacaacgaggattacactatcgttgagcaatatgaaagagc tgaagggcggcactcgacaggtggcatggatgaattgtataag**ggaggaggaggaggtgga***gacaactatgaaagtaag ttctaaaattgtatatgtgttgtttattggttgaaaatgattatttgaattcgctagaaaggaatagtggcgatagcaatcgataaaac aagagagaaactatgcctaaatgtggcctataacatgtgggagttttgtgacaattttgt*

In both fragments: Underlined – 30 bp annealing region for the two fragments (within mCherry coding sequence); Bold – linkers consisting of 6 Glycines (6xGly), positioned before (fragment 1) and after (fragment 2) the mCherry sequence; Italics – f*amk-1* sequence between bp 82 to 248; Sequence in parenthesis (fragment 1) – adapter (adapter was designed to increase G/C content for G-block synthesis).

The annealed product of the two fragments contains an adapter sequence followed by 5’UTR::signal-peptide::6xGly-linker::mCherry::6xGly-linker::N-terminus-famk-1 (bp 82 to 248). The G-block DNA was annealed and amplified using primers specific for the 5’ and 3’ ends and Phusion polymerase (NEB, M0530). Gibson assembly (NEB, E2611) was then used to ligate the PCR product with wild type famk-1 in vector pOD2230/pAG69, thus generating a 5’UTR::signal-peptide::6xGly-linker::mCherry::6xGly-linker::famk-1::famk-1-3’UTR fusion in pOD2318/pAG122.

For expression of ER-retained *famk-1*, KDEL sequence was added at the C-terminus of famk-1, right before the STOP codon, using the following complementary oligos (IDT): Forward – 5’*ggatgataagaaaactgtt<u>aaggatgaacta</u>**tag**tcgtcattcgttaggtcttc*3’ Reverse – 5’*gaagacctaacgaatgacga**cta**<u>tagttcatcctt</u>aacagttttcttatcatcc*3; Both oligos contain a KDEL coding sequence (underlined), followed by a stop codon (bold) and flanked by 19-20 bp homology to *famk-1*. For comparison of WT and ER-retained FAMK-1, Gibson assembly was used ((Gibson et al. 2009), NEB cloning kit, E2611) to insert the above KDEL sequence into pOD2318/pAG122 (mCherry:: famk-1), thus generating a transgene encoding mCherry::famk-1-KDEL (pOD2319/pAG123).

For spermathecal expression of FAMK-1, *sth-1* promoter (p*sth-1*) was used (Bando et al. 2005). A 1380 bp region upstream of the *sth-1* open reading was amplified using: 1-Forward-5’ccgaaagaacggccacaatt3’ and 2-Reverse-5’gttgctctagcacaaaaagac3’ oligos. p*sth-1* was used to drive expression of either *his-72* (transcriptional reporter, pOD2234/pAG81, **Fig. 6 A**), *famk-1* (pOD2240/pAG93, **Fig. 6, B and C**) or *famk-1::gfp* (pOD2243/pAG107, **Fig. S6 B**). In all constructs, a 1761 bp region of the *famk-1* 3’UTR was used (**Fig. 6, A and B** and **Fig. S6 B**). For the transcriptional reporter, Gibson assembly was used to generate p*sth-1::gfp::his-72::6xGly-linker::famk-1-3’UTR* construct ligated into pCFJ151 vector (**Fig. 6 A**).

#### Light microscopy

Images in **Fig. 2, B and C**; **Fig. 5, A, B and D**; **Fig. 6 A**; **Fig. S1 B**; **Fig. S2 A**; **Fig. S4 C**; **Fig. S5**; **Fig. S6 B** were acquired on an inverted Zeiss Axio Observer Z1 system with a Yokogawa spinning-disk confocal head (CSU-X1), a 63x 1.4 NA Plan Apochromat objective (Zeiss, Oberkochen, Germany). Gravid adult worms were dissected and early embryos were collected using mouth pipette. Embryos were mounted on a 2% agarose pad in a 5 µl of M9 buffer, and sealed with a coverslip No.1.5 thickness. 8×1,5 µm z-sections were acquired. Images in **Fig. 2 C** were acquired at 1-minute intervals for 1 hour. Environmental temperature during image acquisition was typically maintained at 19-20°C, except for experiments at which the effect of elevated temperature on phenotype was examined, in which environmental temperature was maintained at 23°C.

Images of furrow ingression in **Fig. S2 A** were acquired at 20 sec intervals. Images in **Fig. 4 E** and **Fig. S2 B** were acquired on a CV1000 spinning disk confocal system (Yokogawa Electric Corporation) with a 512×512 EM-CCD camera (Hamamatsu), 60X 1.35NA U-PlanApo objective and CellVoyager software. Early-stage embryos were mouth pipetted into wells of a glass bottom 384-well Sensoplate (Greiner Bio-One) containing 100 µl of 0.1 mg/ml tetramisole hydrochloride (TMHC) in M9 (to anesthetize worms that later hatch). Before imaging, the plate was spun for 1 minute at 600 x g to settle the embryos. Images were acquired at bright-field and two different wavelengths as follows: (1) bright-field: 90% power, 25 ms, 20% gain (2) 488nm: 100% power, 200 ms, 60% gain (3) 568nm: 45% power, 150 ms, 60% gain). 18×2 µm z-sections were acquired at 20-minute intervals for 10 hours.

In **Fig. 2 A**, to assess the stages of embryonic arrest, the wells were post-scanned in brightfield with 10X objective 20h after embryos were initially mounted. For imaging germlines in **Fig. S1 B** and **Fig. S4 C**, worms were anesthetized by 20-25 min incubation in a mix of 1 mg/ml Tricaine (ethyl 3-aminobenzoate methanesulfonate salt, Sigma-Aldrich, E10521) and 0.1 mg/ml of tetramisole hydrochloride (Sigma-Aldrich, T1512) in M9 before mounting on an agarose pad.

#### Fertility and embryonic lethality analysis

L4 hermaphrodites were singled on plates at 20°C or 23° for 24h (t = 0-24 h). The adult worms were removed to new plates for additional 24h incubation at the same temperatures (t = 24-48 h). This step was repeated once again (t = 48-72 h). For each set of plates, hatched worms and unhatched embryos (embryonic lethal) were counted ~24h hours after the mothers were removed. For the fertility plots, the number of embryos laid during each 24 h interval are presented in **Fig. 1, C and F**, while in other figures, presented are the sum of embryos laid between 0-72 h.

#### Image acquisition of larval defects

To image worms with morphological larval defects compared to wildtype N2 (**Fig. S1 B**), DinoEye USB eyepiece camera was mounted on a dissection scope and Dino-Lite software was used (Version 1.19.2, AnMo Electronics Corporation). Worms are imaged when moving on the plate.

#### Spermathecal overexpression of FAMK-1 by extra-chromosomal arrays

Gonads of 10 gravid adult worms were injected with a DNA mix consisting of the plasmid coding for FAMK-1::GFP driven by STH-1 promoter (pOD2243/pAG107, 50 ng/µl) and a plasmid coding for a pharynx fluorescent marker (pCFJ90 [P*myo-2::mCherry*], 2.5 ng/µl). 4 days after injection, 15 progeny of generation F1 with apparent pharynx marker (which indicates formation of extra-chromosomal arrays) were picked and singled.

After the selected F1 progeny laid eggs, imaging was carried out. Plates with F2 progeny to F1 mothers with apparent FAMK-1::GFP in spermatheca (6/15) were maintained. Brood Size counts were performed for worms of generation F4, and the expression of spermathecal FAMK-1::GFP was confirmed in these.

#### FAMK-1 antibody

FAMK-1 (residues 60-512) was expressed using a modified pI-secSUMOstar vector (LifeSensors) and purified from Hi5 insect cells (Bac-to-Bac, Invitrogen, B85502) (Xiao et al. 2013). The purified protein was sent to Cocalico Biologicals for immunizing rabbits. FAMK-1 specific antibodies were affinity-purified from serum with a HiTrap NHS-activated affinity column (GE Healthcare, 17-0716-01) on which FAMK-1 was immobilized.

#### Immunoblotting

Gravid adult worms were picked, rinsed extensively in M9 buffer and lysed in Loading Buffer (125 mM Tris pH 6.8, 10% glycerol, 2% SDS, 0.71M β-mercaptoethanol, 0.05% Bromophenol blue). Extracts were separated on SDS-PAGE gels and transferred onto nitrocellulose membranes. Immunoblots were blocked in TBST (10 mM Tris, pH 8.0, 150 mM NaCl, 0.1% Tween 20) containing 5% milk followed by overnight incubation at 4°C with the affinity-purified anti-FAMK-1 (diluted 1:1000 in blocking buffer). Immunoblots were washed 6X for 5 min in TBST and incubated with HRP-conjugated anti-rabbit secondary antibody (GE Healthcare, NA934) for 30 min at room temp. After washing as above, the immunoblots were subjected to chemiluminescent substrates and visualized on film.

#### *In vitro* analysis of FAMK-1 substrates

HEK293 cells were grown in Dulbecco’s modified Eagle’s medium (DMEM) with 10% (v/v) fetal bovine serum and 100 µg/ml penicillin/streptomycin (GIBCO, 15140-122) at 37°C with 5% CO_2_. Cells were transfected using Fugene (Roche, E269A) with potential FAMK-1 substrates harboring C-terminal V5-His tags (WT or EE>AA mutants) and either wild type or kinase-defective (D387A) FAMK-1 with a C-terminal FLAG tag. All plasmids used for *in vitro* analysis of FAMK-1 substrates are listed in **Table S2**.

After recovery for 24h, cells were extensively rinsed with Tris-buffered saline (TBS) and were ^32^P-labeled for 6-12h in media consisting of phosphate-free DMEM, 10% FBS (dialyzed) and 0.5 mCi orthophosphate. The cell extracts and media were then harvested for immunoprecipitation. For experiments in which FAMK-1-FLAG::KDEL was co-transfected with Clec-233 V5-His the cells were treated as described above except labeling was carried out in serum-free phosphate-free DMEM containing 0.5 mCi orthophosphate.

*<u>Cell extract immunoprecipitation of FLAG tagged FAMK-1-</u>* Cells were rinsed with TBS and were lysed in 400 µl 50 mM Tris, pH 8.0, 150 mM NaCl, 1.0% NP40 (v/v), 10% glycerol, 0.4 mM EDTA (Modified RIPA buffer) containing protease inhibitors and phosphatase inhibitors. The lysate was cleared by centrifugation at 18,000 x *g* for 10min at 4°C. Protein concentration was determined and equal amounts of protein were added to protein gel loading buffer (extract sample). The remaining extract was pre-cleared using ~30 µl protein G-beads (Thermo Scientific, 20397) and rocking for 15 min at 4°C. The beads were pelleted and the extract was subjected to immunoprecipitation using 25 µl mouse anti-FLAG M2 (Sigma-Aldrich, A2220). Extracts were rocked at 4°C overnight and the beads were pelleted followed by washing 6 times with modified RIPA buffer. The extract sample or FLAG immunoprecipitation from the extract was used in all cases to confirm expression of the transfected construct encoding FAMK-1. To determine the extent of FAMK-1 secretion, FLAG immunoprecipitation of the media was carried out as follows. The media (5 ml) was pre-cleared by centrifugation at 1000 x *g* for 10 min. The supernatant was removed and treated as described above for the cell extract FLAG immunoprecipitations.

*<u>Media immunoprecipitation of V5 tagged lectins-</u>* Cell debris was removed by centrifugation at 1000 x *g* for 10 min and the media supernatant then pre-cleared by addition of protein G-beads as above. After removal of the beads, the media was immunoprecipitated using 1 µl rabbit anti-V5 antibody (Millipore, AB3792) and 25 µl of protein G-beads rocked at 4°C overnight. The beads were collected and washed as described for the FLAG immunoprecipitations. The V5 immunoprecipitates were analyzed by immunoblotting and autoradiography.

For CLEC-110, which was not secreted efficiently into the medium, V5 immunoprecipitation from cell extracts was necessary; in this case, the extract was first used for immunoprecipitation with rabbit anti-V5 antibody, and then immunoprecipitated as above using anti-FLAG M2.

#### Phylogenetic analysis

Amino acid sequences of the FAMK-1 orthologues were aligned using Clustal O (Sievers et al. 2011) and the tree was generated using free software phylodendron (iubio.bio.indiana.edu/treeapp/treeprint-form.html).

#### Calcium signaling in the spermatheca during embryo transits

*<u>Image acquisition-</u>*Animals were egg prepped by dissolving gravid hermaphrodites in a solution of sodium hydroxide and sodium hypochlorite and washing the remaining eggs 3x with M9 buffer (reviewed in (Stiernagle 2006)). Drops of M9 containing eggs were plated onto NGM plates seeded with *E. coli* OP50, and raised at 20° C for 68 hours. Animals close to their first ovulation were selected for imaging, anaesthetized with 0.01% tetramisole and 0.1% tricaine in M9 buffer, and mounted on 2% agarose pads. Images were acquired on a Nikon Eclipse 80i microscope, using a 60x, 1.40 NA objective. Fluorescence was monitored using a standard GFP filter set (EX 470/40, BS 495, EM 525/50) (Chroma Technology; Bellows Falls, VT, USA), with fluorescence excitation provided by a Nikon Intensilight CHGFI 130W mercury lamp, attenuated at the source with the ND16 neutral density filter, and shuttered between exposures using a SmartShutter (Sutter Instruments; Novato, CA, USA) triggered by the camera TTL signal. Images were captured using a SPOT RT3 monochrome CCD camera and acquisitions were controlled by SPOT Advanced 5 software (Diagnostic Instruments; Sterling Heights, MI, USA). Images were acquired at a rate of 1 frame per second, an exposure time of 75 ms, and a gain of 8. The full chip of the camera was used, generating images 1600 x 1200 pixels in size. Time-lapse image sequences were saved as 8-bit TIFF files.

*<u>Image processing-</u>* All image processing was done using Fiji (http://fiji.sc/). Each time-lapse image sequence was registered to a single frame, rotated to put the spermatheca in a standard orientation, and cropped to a standard size of 800 x 400 pixels (100 µm x 50 µm). Following this standardization step, GCaMP time series values (F) were generated for each frame of the image sequence by calculating the mean pixel intensity (total pixel intensity / area). For each time series, the baseline GCaMP value (F0) was obtained by averaging the 30 frames prior to oocyte entry, and the normalized GCaMP time series (F/F0) was calculated by dividing the entire GCaMP time series by the baseline GCaMP value. Normalized GCaMP time series are shown plotted in GraphPad Prism (GraphPad Software; La Jolla, CA, USA).

Heat maps of multiple GCaMP time series were generated using Matlab (The MathWorks; Natick, MA, USA). Each time series was cropped to start at the 50^th^ frame prior to oocyte entry and was appended with zeros until the length matched the longest time series in the set. The time series were then assembled into a matrix where each row is a different time series and time increases with the columns. This matrix was then displayed as an image and the color map was scaled so all heat maps were displaying the same range.

---

**Table S1.**
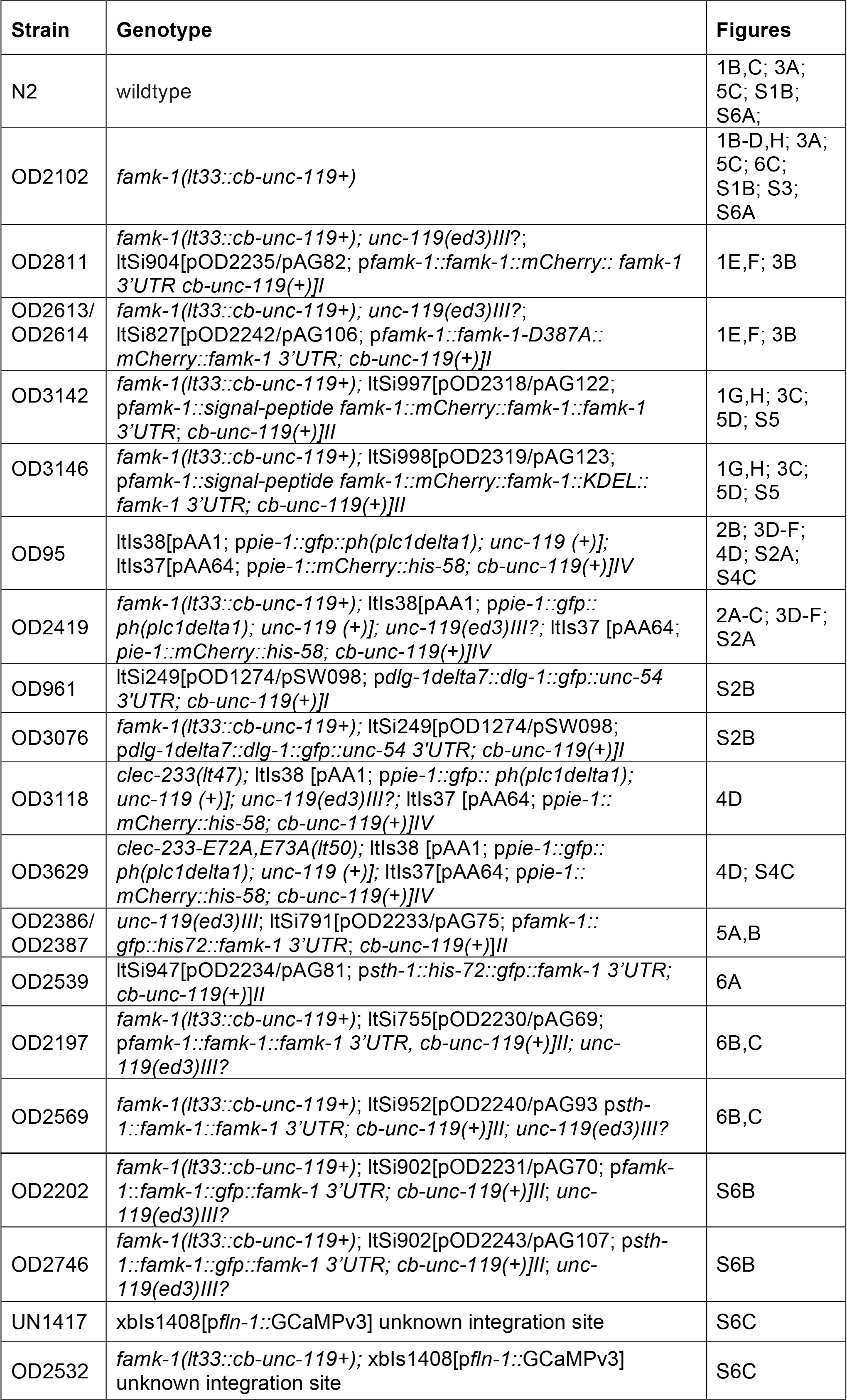
*C. elegans* strains used in this study Strain.

**Table S2.**
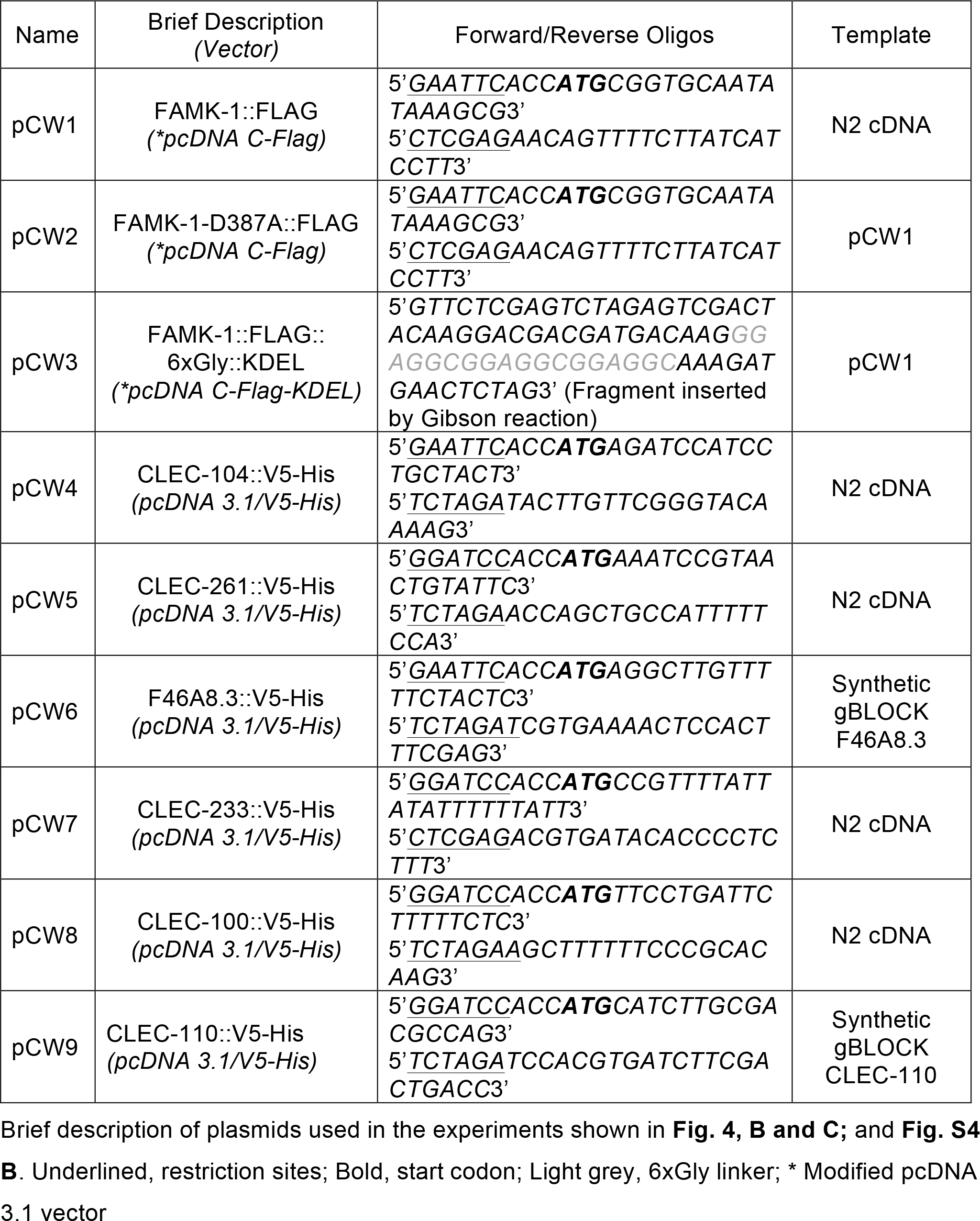
Plasmids used for *in vitro* analysis of FAMK-1 substrates.

**Table S3:** List of hits from proteome-wide "SSSEE" motif search and which are predicted to have a signal peptide and/or transmembrane domain(s)

| <u>Sequence Name</u> | <u>Gene Name</u> | <u>Sequence Name</u> | <u>Gene Name</u> |
| --- | --- | --- | --- |
| ZK666.5 | clec-59 | C06E1.10 | rha-2 |
| Y53H1A.3 | clec-100 | C13G5.2 |  |
| Y18D10A.10 | clec-104 | F31D5.5 | mth-1 |
| Y18D10A.12 | clec-106 | F41E6.12 |  |
| F17B5.3 | clec-109 | F45F2.7 | otpl-7 |
| F17B5.5 | clec-110 | T04C4.1 |  |
| Y116A8A.3 | clec-193 | T14A8.2 |  |
| F21H7.4 | clec-233 | ZK355.5 | irld-65 |
| ZC15.6 | clec-261 | ZC101.2 | unc-52 |
| F49F1.11 |  | F30A10.8 | stn-1 |
| F46A8.3 |  | ZK897.1 | unc-31 |
| F46A8.5 |  | F58E10.1 | ric-7 |
| F46A8.8 |  |  |  |
| K11G12.7 | acr-3 |  |  |
| ZK896.8 | gcy-18 |  |  |
| F35D11.2 | pqn-35 |  |  |
| C33G3.5 |  |  |  |
| C39B10.2 | lgc-41 |  |  |
| F09B9.4 |  |  |  |
| F26C11.3 |  |  |  |
| F53F8.4 |  |  |  |
| H19J13.1 | otpl-8 |  |  |
| R05A10.2 |  |  |  |
| R06F6.7 |  |  |  |
| T06D8.5 | cox-15 |  |  |
| T07D10.1 |  |  |  |
| T28D6.5 |  |  |  |
| W03G11.4 |  |  |  |
| Y51A2A.6 |  |  |  |
| Y65A5A.2 |  |  |  |
| B0207.10 |  |  |  |

**Figure S1.**
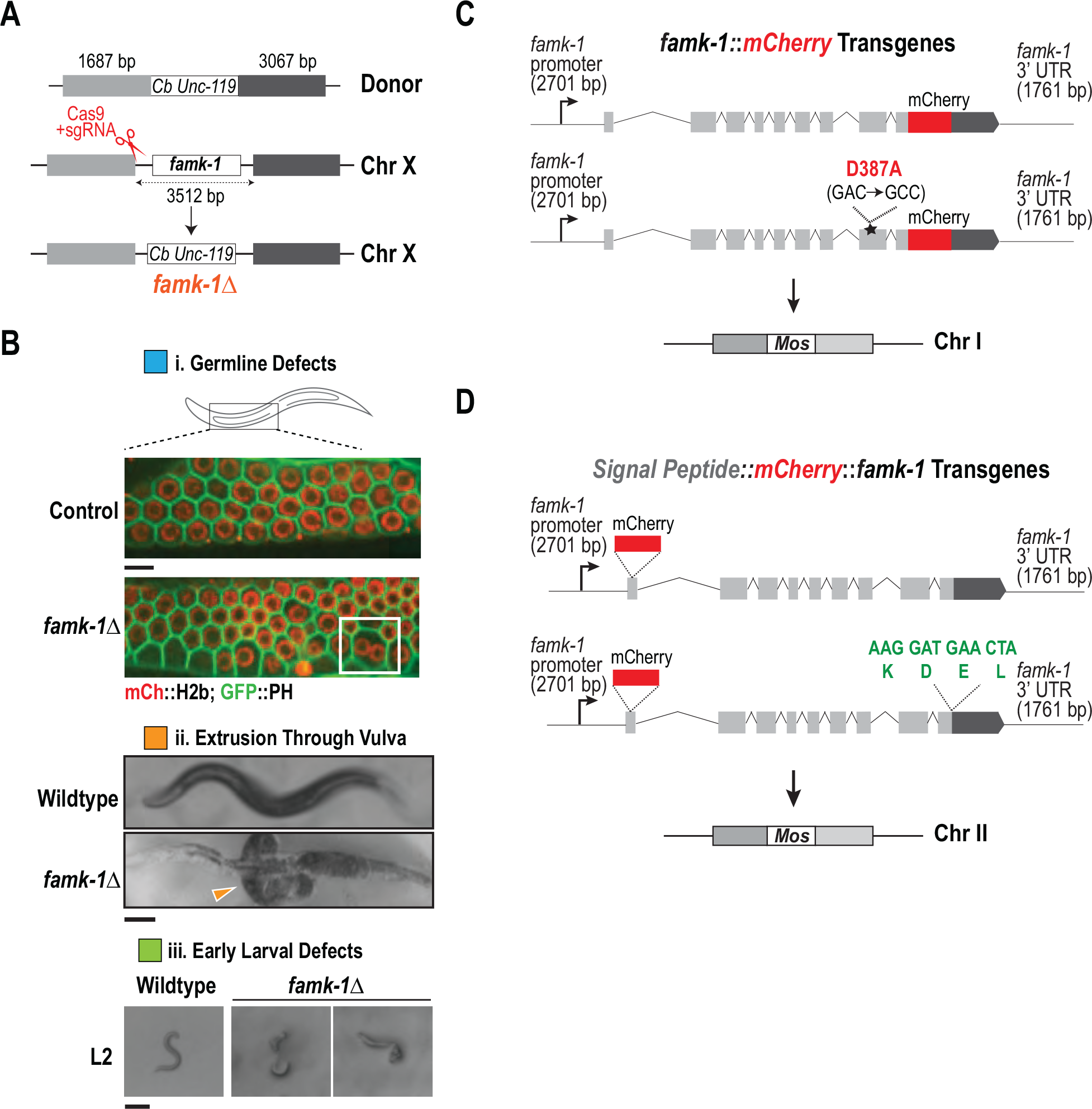
Generation of the *famk-1*Δ mutant, *famk-1* single copy transgenes, and images of phenotypes observed in *famk-1*Δ. **(A)** Schematic of *famk-1*Δ mutant generation. The entire *famk-1* coding sequence was replaced with the *Cb unc-119* marker. **(B)** Images of post-hatching defects observed in *famk-1*Δ larvae and adults. Scale bar in B(i), 5 µm; Scale bars in B(ii) & B(iii), 100 µm. **(C)** Schematic of *famk-1::mCherry* transgene. **(D)** Schematic of wildtype and KDEL *mCherry::famk-1* transgenes.

**Figure S2.**
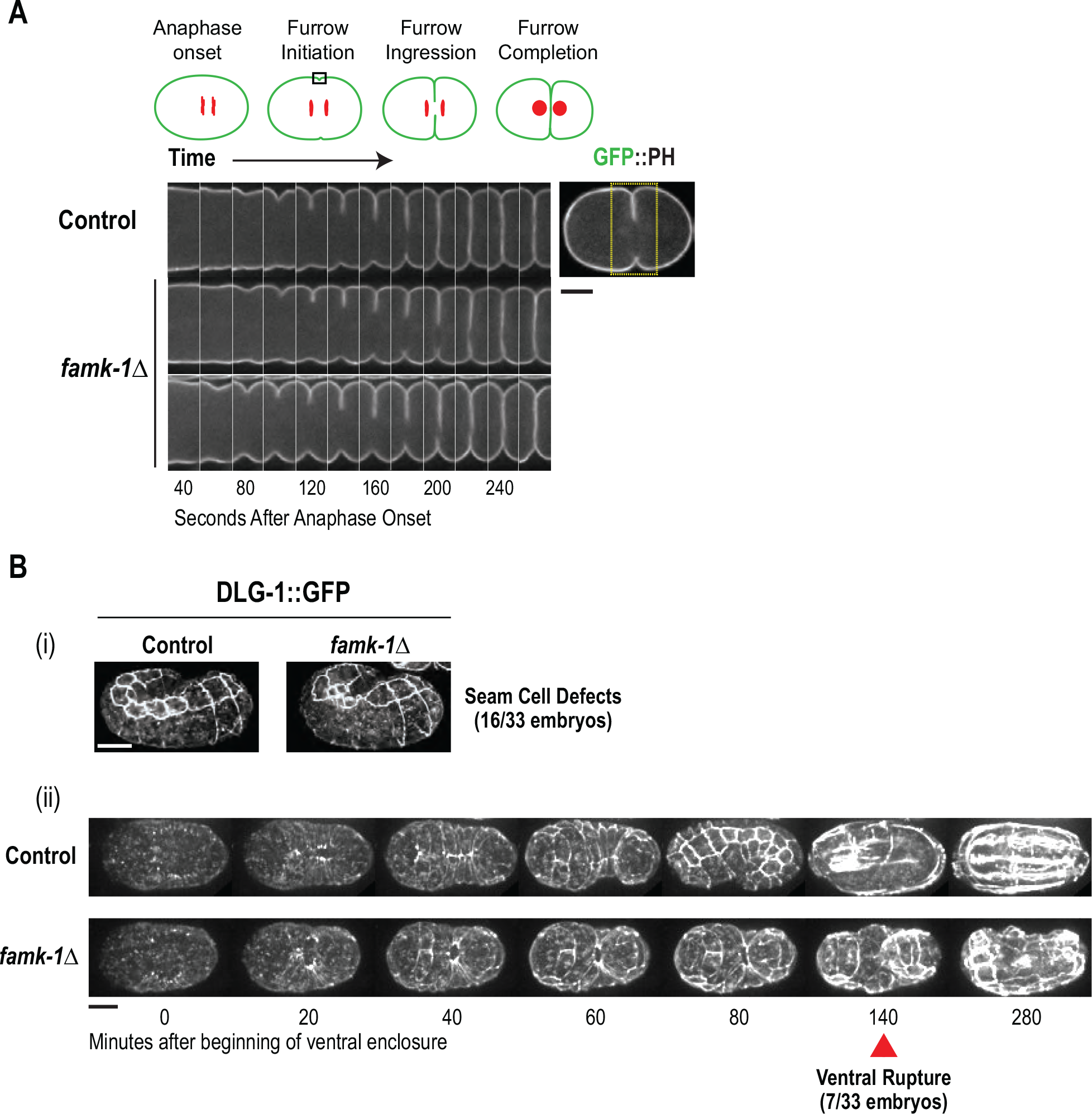
Embryonic phenotype analysis of *famk-1*Δ. **(A)** One-cell embryo cytokinesis imaging of control and *famk-1*Δ embryos expressing GFP-fused plasma membrane marker. Similar results were observed in 10 imaged control and *famk-1*Δ embryos. Scale bar, 10 µm. **(B)** Imaging of DLG-1::GFP in late stage control and *famk-1*Δ embryos. Scale bar, 10 µm.

**Figure S3.**
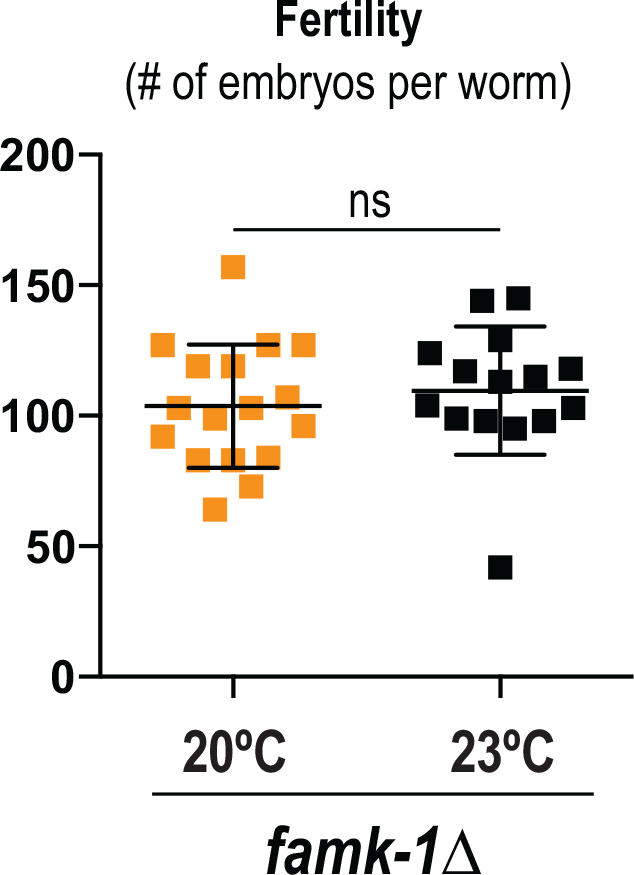
Analysis of *famk-1*Δ fertility at 20°C and 23°C.

**Figure S4.**
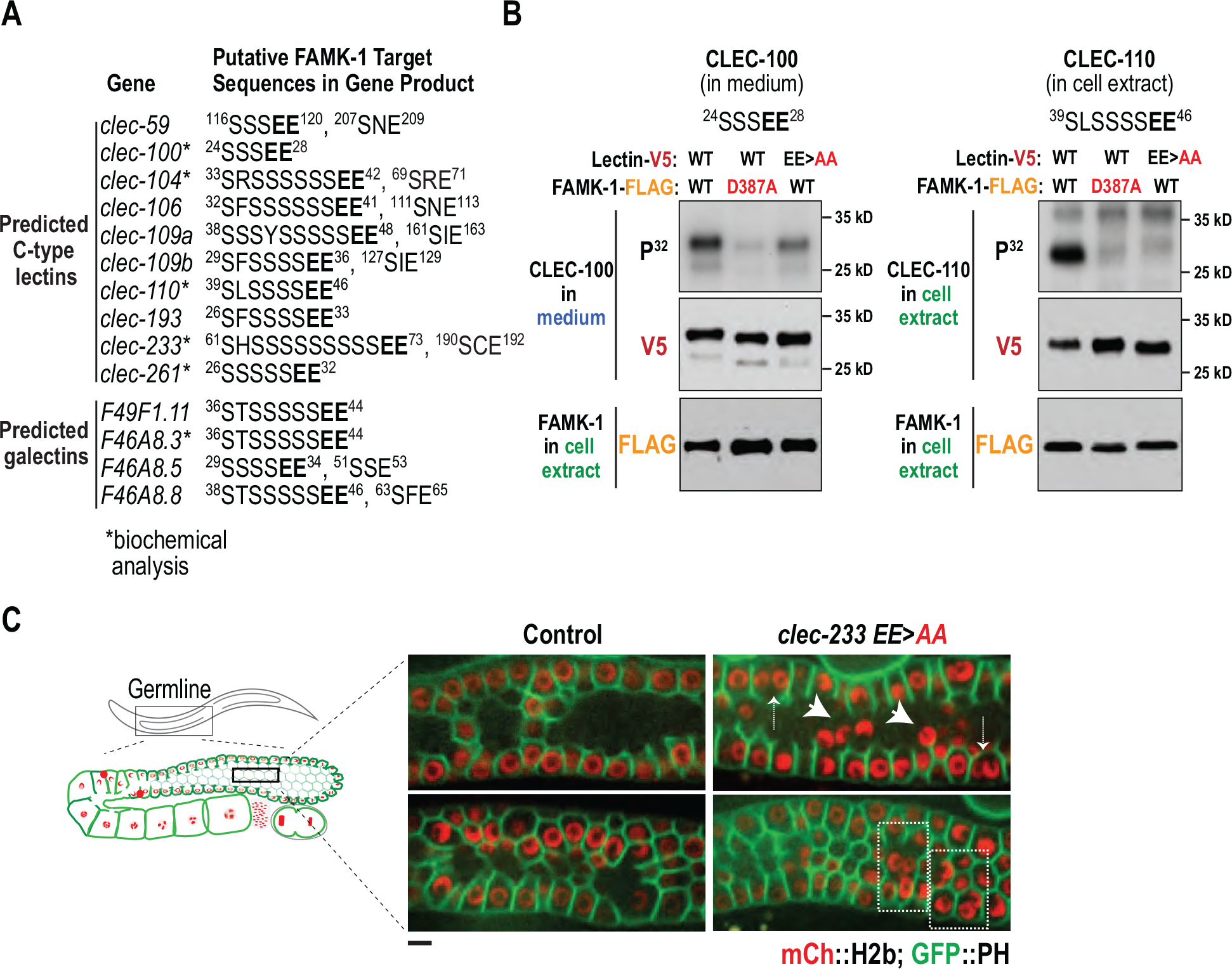
Analysis of lectins as putative substrates of FAMK-1. **(A)** Motifs identified by the proteome-wide search for ‘SSSEE’ motifs. Additional ‘SxE’ motifs in the same target are also listed. Residue numbers are indicated to identify location within the primary sequence. The products of genes marked with asterisks were subjected to biochemical analysis after co-expression with FAMK-1 in HEK293 cells (see *Fig. 4 B*). **(B)** Analysis of CLEC-100 and CLEC-110 phosphorylation by FAMK-1. CLEC-110 was not efficiently secreted and thus its phosphorylation status was analyzed in cell extracts rather than in the medium. **(C)** Images of the germline in control and *clec-233 EE>AA* worms expressing fluorescent fusions that label the plasma membrane and chromatin. Arrows highlight nuclei delaminating from the gonad surface and boxes delineate regions with multinucleation. Similar defects were observed in 3/10 imaged *clec-233 EE>AA* worms. Scale bar, 5 µm.

**Figure S5.**
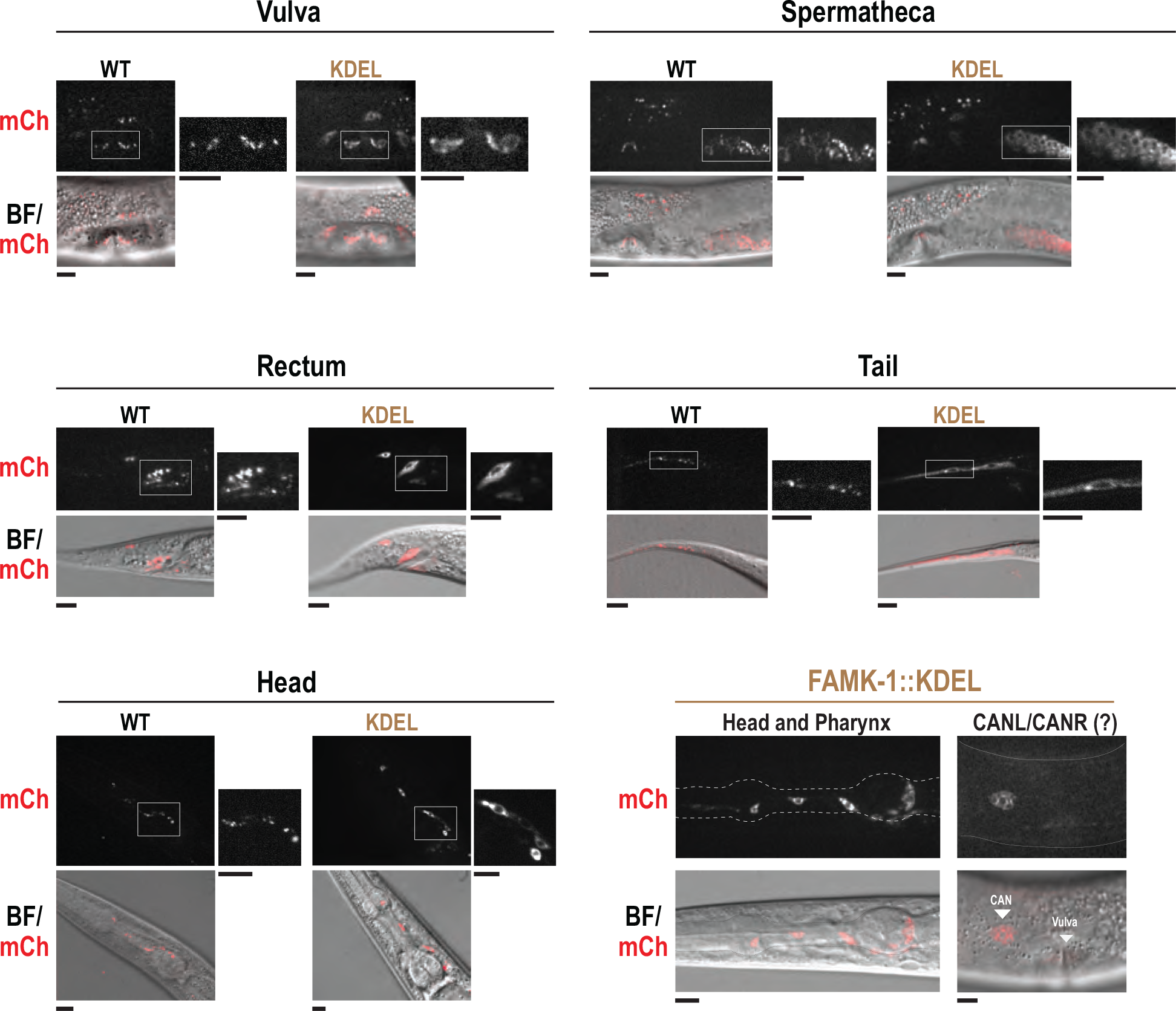
Images of WT and KDEL mCherry::FAMK-1 in different tissues. mCherry signal and overlay of brightfield (BF) and mCherry images are shown. Representative images are shown for each tissue; at least 6 worms were imaged per tissue. Scale bars, 10 µm.

**Figure S6.**
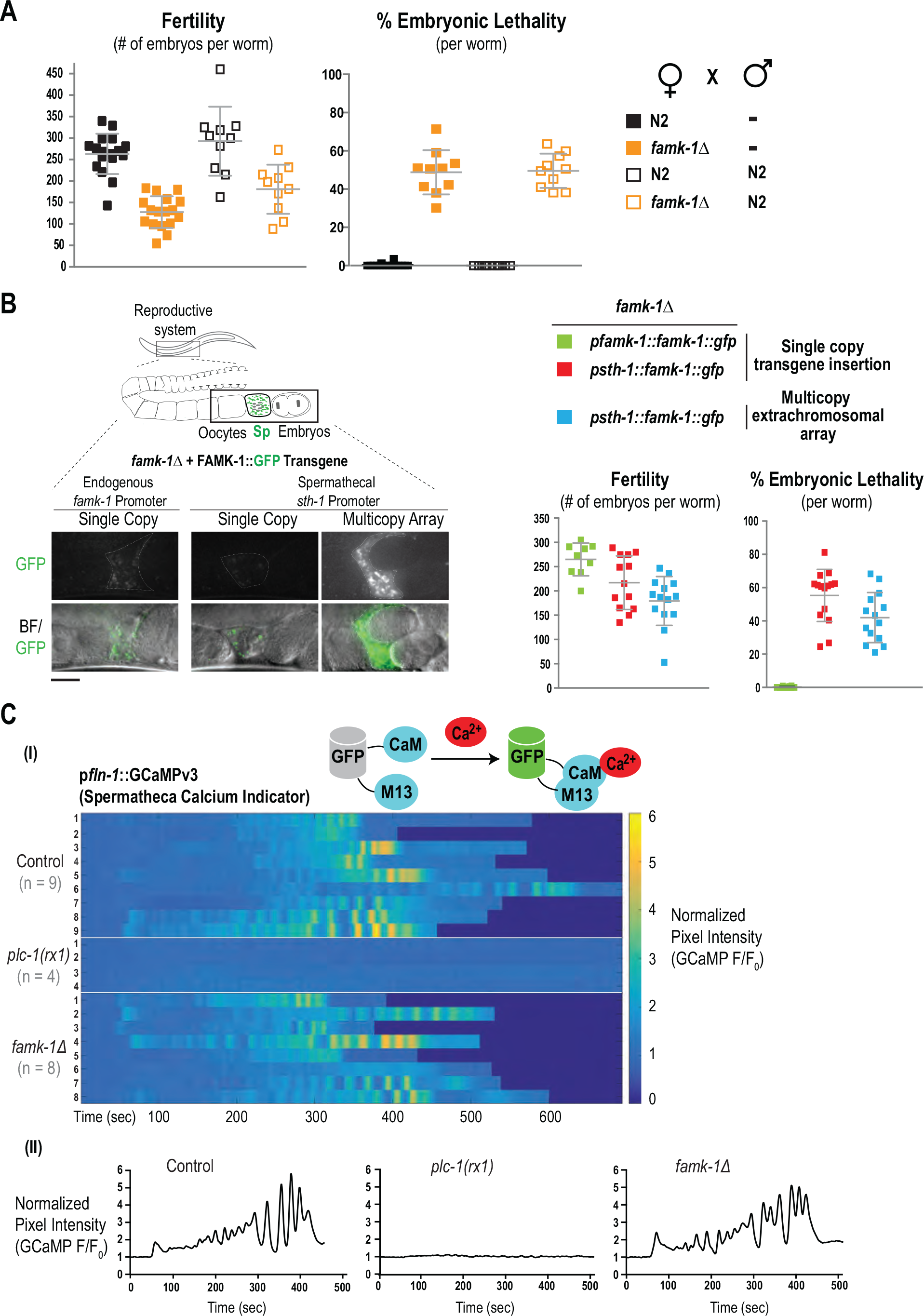
Analysis of spermatheca-expressed FAMK-1. **(A)** Mating with N2 males does not rescue the fertility defect or embryonic lethality of *famk-1*Δ. Successful mating was confirmed by presence of males among the F1 progeny. **(B)** *(left)* lmages of FAMK-1::GFP expressed under control of the spermathecal promoter p*sth-1* (single copy or multicopy array) or under control of the endogenous promoter, p*famk-1* (single copy). Overlay images of brightfield (BF) and GFP signal images are shown below. Scale bar, 20 µm. *(right)* Effect of overexpression of FAMK-1 from an extrachromosomal array on fertility and embryonic viability at 20°C. **(C)** Analysis of calcium oscillations in the spermatheca using p*fln-1*::GCaMPv3 during the first few ovulations of *C. elegans*. Each row represents an individual imaged worm. *plc-1(rx1)* serves as a negative control, in that calcium oscillations are not observed in the spermatheca in this mutant. Plots below show the normalized pixel intensity for representative individual worms.

